# Mesoscale regulation of MTOCs by the E3 ligase TRIM37

**DOI:** 10.1101/2024.10.09.617407

**Authors:** Zhong Y. Yeow, Sonia Sarju, Mark v. Breugel, Andrew J. Holland

## Abstract

Centrosomes ensure accurate chromosome segregation during cell division. Although the regulation of centrosome number is well-established, less is known about the suppression of non-centrosomal MTOCs (ncMTOCs). The E3 ligase TRIM37, implicated in Mulibrey nanism and 17q23-amplified cancers, has emerged as a key regulator of both centrosomes and ncMTOCs. Yet, the mechanism by which TRIM37 achieves enzymatic activation to target these mesoscale structures had remained unknown. Here, we elucidate TRIM37’s activation process, beginning with TRAF domain-directed substrate recognition, progressing through B-box domain-mediated oligomerization, and culminating in RING domain dimerization. Using optogenetics, we demonstrate that TRIM37’s E3 activity is directly coupled to the assembly state of its substrates, activating only when centrosomal proteins cluster into higher-order assemblies resembling MTOCs. This regulatory framework provides a mechanistic basis for understanding TRIM37-driven pathologies and, by echoing TRIM5’s restriction of the HIV capsid, unveils a conserved activation blueprint among TRIM proteins for controlling mesoscale assembly turnover.

## Main

Mesoscale protein assemblies serve as organizational hubs that dictate the spatial arrangement of subcellular components. The centrosome is one prominent example, serving as the primary microtubule-organizing center (MTOC) in animal cells that orchestrates the accurate segregation of chromosomes during cell division^1^. Centrosomes consist of a pair of centrioles nestled within a proteinaceous matrix known as the pericentriolar material (PCM). The PCM is an assembly of several hundred proteins that collectively act to anchor and nucleate microtubules^2,3^. Recent advancements in super-resolution microscopy have shown that the interphase PCM comprises an organized assembly of radial protein layers surrounding the centriole^4^.

Centrosome number is rigorously controlled in tandem with dynamic changes in the PCM’s composition and volume as cells progress through the cell cycle^5,6^. This intricate regulation underpins the centrosome’s crucial function in cell division, where numerical aberrations can give rise to a range of pathologies, including cancer and neurodevelopmental disorders^7,8^. Some differentiated cell types utilize non-centrosomal MTOCs (ncMTOCs) in interphase for specialized functions^9^. Crucially, the presence of ncMTOCs during mitosis can threaten genome integrity^10,11^, but the regulatory mechanisms governing their formation remain poorly understood^9^.

TRIM37 is a member of the TRIpartite Motif (TRIM) family of proteins characterized by the conserved RBCC motif, which includes a <u>R</u>ING E3 ubiquitin ligase domain, a <u>B</u>-box domain, and a <u>C</u>oiled-<u>c</u>oiled domain^12^. Loss-of-function mutations in *TRIM37* cause Mulibrey nanism (MUL), a rare autosomal recessive disorder characterized by growth failure and multi-organ abnormalities^13^. Initial reports of TRIM37’s localization to peroxisomes led to the classification of MUL as a peroxisomal disorder^14^. However, TRIM37-deficient mice do not display peroxisome abnormalities despite recapitulating key features of the human disease^15^. Recent work has recast TRIM37 as a central player in centrosome regulation^16–18^. In MUL patient fibroblasts, loss of TRIM37 causes the centriolar protein Centrobin to accumulate as a single highly structured cytoplasmic assembly. These assemblies emerge and detach from the centrosome to act as ncMTOCs that promote chromosome segregation defects–a process now implicated as a key driver of MUL pathogenesis^19,20^.

While *TRIM37* loss-of-function mutations cause MUL, TRIM37 overexpression frequently occurs during tumorigenesis^21^. *TRIM37* is located within 17q23, a chromosome region often amplified in breast cancer or gained in neuroblastomas^22,23^. Amplicon-directed overexpression of TRIM37 promotes the degradation of the PCM scaffolding protein CEP192, leading to reduced PCM levels at the centrosome and an increased frequency of mitotic errors^24^. Moreover, cancer cells exhibiting elevated TRIM37 expression are therapeutically vulnerable to centrosome loss induced by PLK4 inhibition. This arises as high levels of TRIM37 impede the formation of CEP192-containing foci, a ncMTOC crucial for mitotic spindle assembly in cells lacking centrosomes^24,25^. The discovery of this synthetic lethal interaction has spurred the development of PLK4 inhibitors, now entering clinical trials to target tumors overexpressing TRIM37.

TRIM37 has emerged as a critical regulator of centrosome function that counteracts the formation of ectopic Centrobin assemblies and degrades the PCM scaffolding protein CEP192, but how TRIM37 recognizes its substrates in the form of mesoscale cellular assemblies remains unclear. Here, we demonstrate that substrate assembly promotes TRIM37 oligomerization, a pivotal step that activates its ubiquitin ligase function. This activation mechanism enables the selective degradation of centrosome proteins incorporated into higher-order assemblies, providing an elegant solution through which TRIM37 exerts control over cellular structures integral to cell division.

## Results

### Mulibrey Nanism (MUL) mutations in *TRIM37* reveal a common framework for the regulation of centrosomes and non-centrosomal Centrobin assemblies

TRIM37 possesses a core RBCC motif, followed by a unique TRAF domain and an unstructured C-terminal tail (Fig. 1b, center)^26^. To examine the contributions of these domains to TRIM37 function, we knocked out *TRIM37* in non-transformed RPE-1 and re-expressed wild-type (WT) or mutant variants of HA-tagged TRIM37. Knockout of *TRIM37* was confirmed by Sanger sequencing and loss of TRIM37 protein expression (Extended Data Fig. 1a, b).

**Figure 1.**
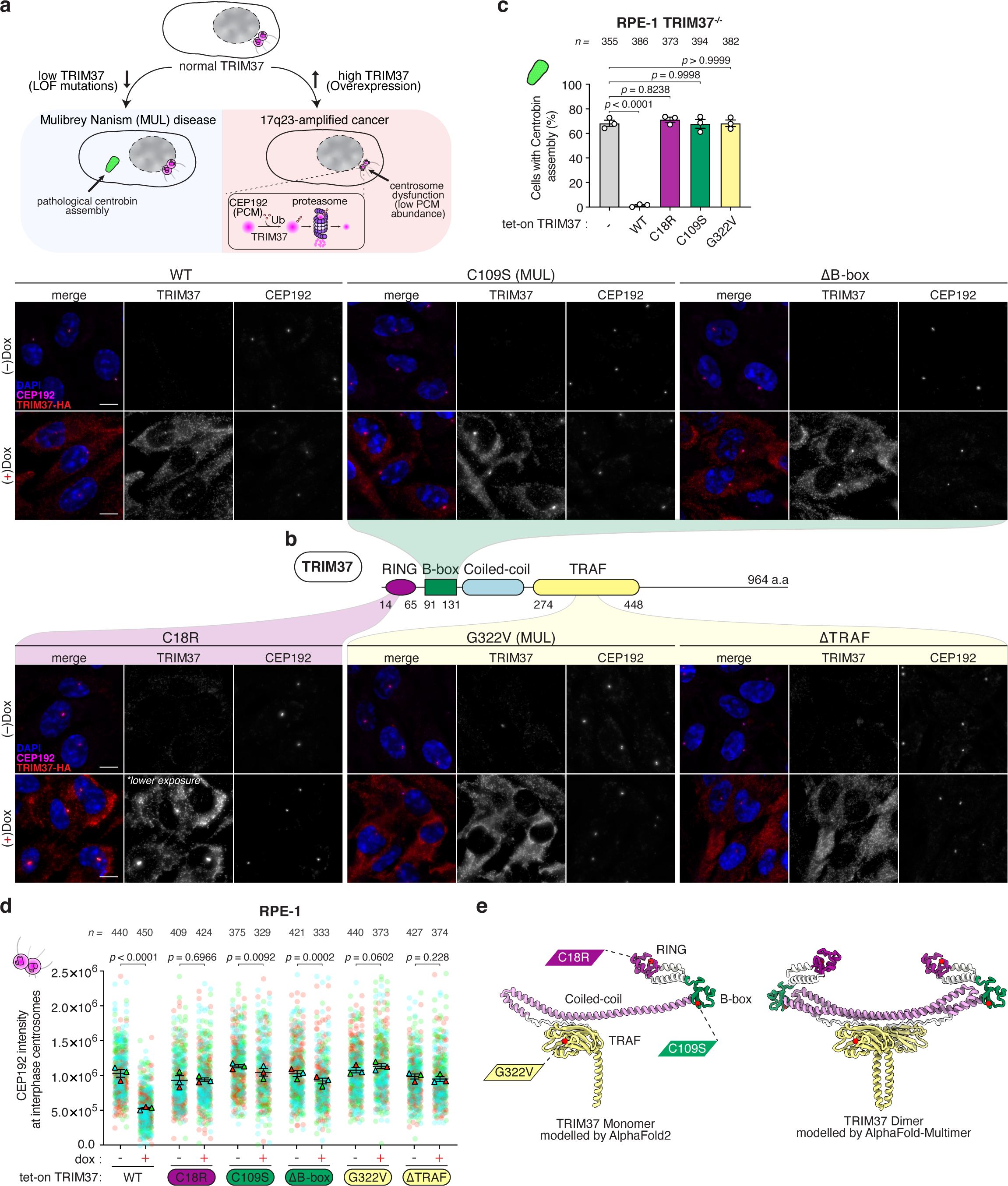
Domain-specific impact of Mulibrey nanism (MUL) *TRIM37* mutations on MTOC regulation. (A) A diagram depicting TRIM37 dysregulation in MUL and 17q23-amplified cancers. In MUL, TRIM37 loss-of-function mutations result in the aberrant formation of Centrobin assemblies, which act as ectopic MTOCs during mitosis. Conversely, in 17q23-amplified cancers, elevated expression of TRIM37 leads to excessive degradation of centrosomal CEP192. (B) Centre, domain architecture of TRIM37 (UniProt ID 094972) highlighting the common RBCC motif (RING, B-box, Coiled-coil domains) and unique C-terminal TRAF domain. Surrounding panels, localization pattern and effect of HA-tagged TRIM37 variants (domain-specific mutations and deletions) on centrosomal CEP192 levels in RPE-1 tet-on TRIM37 cells. MUL indicates Mulibrey nanism patient-derived mutations. Representative data; *n* = 3 biological replicates. Scale bars, 5 μm. (C) Quantification of Centrobin assembly occurrence in RPE-1 *TRIM37*^-/-^ cells expressing the indicated HA-tagged TRIM37 variants. *n* = 3 biological replicates, each with >100 cells. *P* values, one-way ANOVA with post hoc Dunnett’s multiple comparisons test to evaluate differences between the TRIM37 variants and wild-type (WT). Mean ± s.e.m. (D) Quantification of centrosomal CEP192 signal in RPE-1 tet-on TRIM37 cells from (B). *n* = 3 biological replicates, each with >100 cells. *P* values, unpaired two-tailed *t*-test. Mean ± s.e.m. (E) Left, AlphaFold-predicted monomer of TRIM37. The RING, B-box, Coiled-coil, and TRAF domains are shown, with mutated residues highlighted in red. Right, AlphaFold Multimer-predicted model of a TRIM37 dimer. For both models, the unstructured C-terminal tail of TRIM37 (residues 449–964) is not shown due to the lack of a high-confidence prediction.

Consistent with prior reports^19,20^, *TRIM37*^-/-^ cells formed cytoplasmic Centrobin assemblies (Fig. 1a and Extended Data Fig. 1c, left panel) that were lost upon the expression of WT TRIM37 (Fig. 1c and Extended Data Fig. 1c). Inactivation of TRIM37 E3 ligase activity with the (C18R) RING domain mutation^19,24,25^ prevented degradation of the Centrobin assembly without impacting the recruitment of TRIM37 to the assembly (Fig. 1c and Extended Data Fig. 1c).

Clinically relevant MUL mutations within the B-box (C109S)^27^ and TRAF domain (G322V)^28^ were also defective in degrading the Centrobin assembly, supporting a causative role of this assembly in MUL pathogenesis (Fig. 1c). Notably, while the (C109S) B-box mutant localized to the Centrobin assembly, the (G322V) TRAF mutant did not (Extended Data Fig. 1c).

To assess the relevance of these findings in the context of 17q23-amplified cancers, we monitored the impact of doxycycline-induced overexpression of TRIM37 on the abundance of its centrosomal substrate CEP192 (Fig. 1a). Expression of WT TRIM37 in RPE-1 cells drove a significant reduction in CEP192 levels at the centrosome (Fig. 1b,d). In contrast, TRIM37 C18R, C109S, and G322V mutants were ineffective at degrading centrosomal CEP192 (Fig. 1b,d).

Reflecting the localization patterns seen with the Centrobin assembly, both the TRIM37 C18R RING and C109S B-box mutants localized to the centrosome, while the G322V TRAF mutant failed to do so (Fig. 1b). Deletion of the B-box or TRAF domain (ΔB-box and ΔTRAF) phenocopied the effects of the respective MUL point mutants (Fig. 1b,d and Extended Data Fig. 1d,e), indicating that these mutations lead to domain-specific loss-of-function in TRIM37.

Collectively, these data suggest that TRIM37 employs a common mechanism for the recognition and subsequent degradation of Centrobin in cytoplasmic assemblies and CEP192 incorporated into centrosomes.

### The TRIM37 TRAF domain plays a central role in centrosomal substrate recognition

The TRIM family member TRIM5 is known for its role in inhibiting retroviral infections, particularly HIV^29^. TRIM5 and TRIM37 have a similar domain organization, except the TRAF domain of TRIM37 is replaced by a SPRY domain in TRIM5. TRIM5 assembles into a dimer, with the two SPRY domains centrally located and each monomer’s RING and B-box domains positioned at opposite ends of an antiparallel coiled-coil^30,31^. As RING dimerization is crucial for E3 ligase activity, this antiparallel configuration prevents the interaction of the two RING domains within a single TRIM5 dimer. E3 activation occurs when many TRIM5 dimers bind to the surface of the viral capsid through the SPRY domain and assemble into an oligomeric lattice. This facilitates the dimerization of RING domains from adjacent TRIM5 dimers and subsequent E3 ligase activity^32,33^. The crystal structure of the TRIM37 RING dimer closely resembles that of TRIM5^33^. Moreover, AlphaFold2 modeling^34^ showed a high-confidence dimer prediction for TRIM37 that was similar to TRIM5, with the TRIM37 TRAF domain occupying the position of the TRIM5 SPRY domain (Fig. 1e). Given that mutation or deletion of the TRIM37 TRAF domain prevented recruitment of TRIM37 to Centrobin assemblies and the centrosome, the TRAF domain is likely to be the substrate recognition motif of TRIM37.

To identify TRAF domain-mediated interactors of TRIM37, we performed proximity-dependent biotin labeling with mTurbo-tagged TRIM37 (Fig. 2a,b). We hypothesized that the (C18R) RING mutant would show extensive labeling of centrosome substrates due to its impaired ability to promote substrate degradation, while the (G322V) TRAF mutant would exhibit a reduced labeling profile. We curated a list of 98 high-confidence TRAF-mediated TRIM37 proximity-interaction partners after background subtraction (Fig. 2c). Of these interactors, 74% (73/98) overlapped with published centrosome proximity datasets^35,36^, with CEP192 and Centrobin among the most enriched proteins (Fig. 2d). Gene ontology analysis further emphasized the significant enrichment of centrosome and ciliary proteins within the TRIM37 proximity interactome (Fig. 2e), underscoring the central role of the TRIM37 TRAF domain in centrosomal substrate targeting.

**Figure 2.**
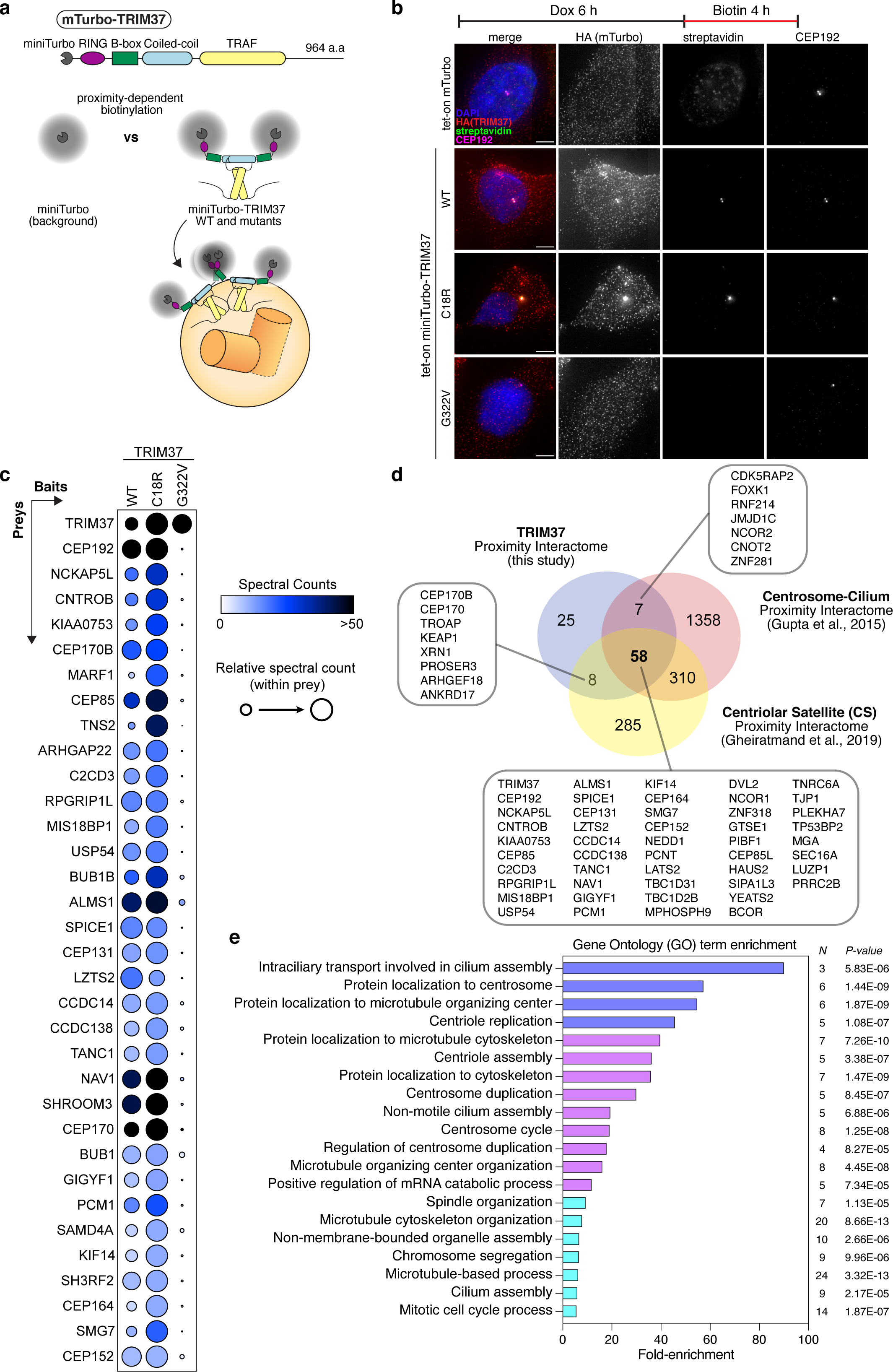
Proximity-dependent biotin identification (BioID) identifies TRAF domain interactors of TRIM37. (A) Top, schematic of miniTurbo-TRIM37 construct used for BioID labelling experiments. Bottom, depiction of the approach to isolate TRAF domain-specific interactors of TRIM37. (B) Immunofluorescence images of biotin-labelled proteins in RPE-1 cells expressing the indicated mTurbo constructs. Streptavidin staining indicates biotinylated proteins, with centrosomes marked by CEP192 staining. Representative data; *n* = 2 biological replicates. Scale bars, 5 μm. (C) Thresholded mass spectrometry results displaying the top 34 proximity interactors (TRAF-domain specific) by spectral count. Interactors were identified using filters detailed in the Methods section, highlighting differential labelling by TRIM37 mutants after background subtraction. (D) Venn diagram illustrating the overlap between TRIM37 TRAF domain-specific proximity interactome (this study) and two published centrosome proximity interactomes. Accompanying list specifies hits common to the TRAF-domain interactome. (E) Gene ontology analysis of mass spectrometry data from BioID experiments.

### SPRY-TRAF domain swap repurposes HIV restriction factor TRIM5 as a MTOC regulator

If TRIM5 and TRIM37 share structural and regulatory principles, we reasoned we could impart the regulation of TRIM37 substrates by swapping the TRIM5 SPRY domain with the TRIM37 TRAF domain (Fig. 3a). TRIM5 did not localize to the centrosome and could not degrade CEP192 (Fig. 3b–d). By contrast, the TRIM5-TRAF chimera localized to the centrosome and degraded CEP192 to a similar extent as TRIM37 (Fig. 3b–d). Moreover, introducing the MUL TRAF domain mutation (G322V) into the TRIM5-TRAF chimera abolished its centrosomal localization and capacity to degrade CEP192 (Fig. 3b–d). Similar findings were observed with the degradation of the Centrobin assembly in *TRIM37^-/-^* cells (Fig. 3e,f). These data support the proposal that TRIM37 functions analogously to TRIM5, with the TRAF domain being key for the selective regulation of MTOCs.

**Figure 3.**
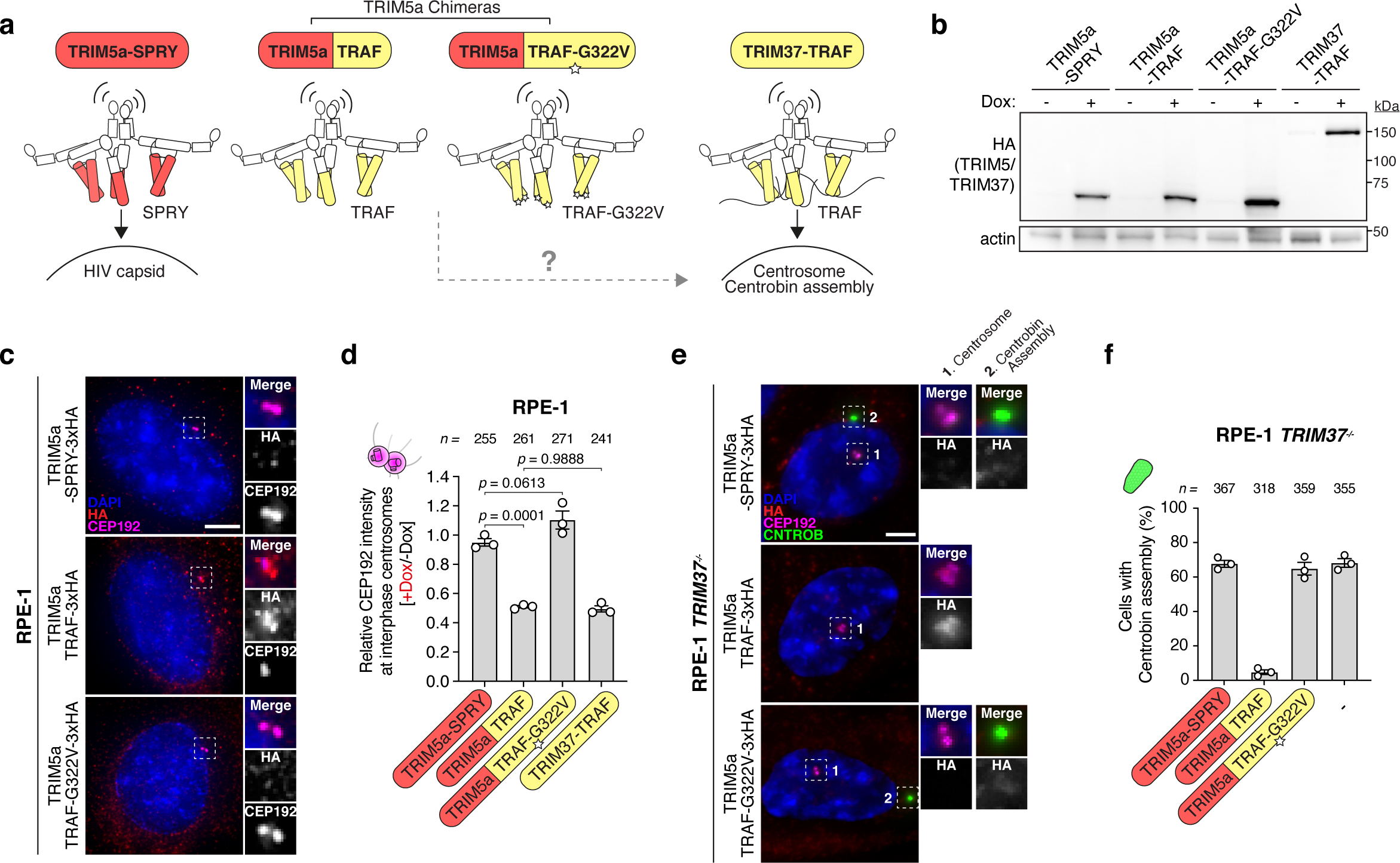
Chimeric TRIM5 bearing the TRIM37 TRAF domain regulates MTOCs. (A) Schematic overview of the domain swap strategy, which replaces the TRIM5 SPRY domain with the TRIM37 TRAF domain to generate a chimeric TRIM5-TRAF protein. (B) Immunoblot showing total protein expression levels of indicated HA-tagged TRIM5 constructs in RPE-1 tet-on TRIM5 cells. Actin, loading control. Representative data; *n* = 3 biological replicates. (C) Representative images of the localization and effect of indicated HA-tagged TRIM5 constructs on centrosomal CEP192 levels in RPE-1 tet-on TRIM5 cells. Representative data; *n* = 3 biological replicates. Scale bars, 5 μm. (D) Quantification of centrosomal CEP192 signal upon doxycycline-induced expression of indicated constructs in RPE-1 tet-on TRIM5 cells from (C), with TRIM37 included as a benchmark. *n* = 3 biological replicates, each with >100 cells. *P* values, one-way ANOVA with post hoc Tukey’s multiple comparisons test. Mean ± s.e.m. (E) Representative images of RPE-1 *TRIM37*^−/−^ cells expressing the indicated HA-tagged TRIM5 constructs. Inset #1 denotes the centrosome, marked by CEP192, and inset #2 denotes the Centrobin assembly, identified by intense Centrobin staining that is non-centrosome localized. Representative data; *n* = 3 biological replicates. Scale bars, 5 μm. (F) Quantification of Centrobin assembly occurrence in RPE-1 *TRIM37*^-/-^ cells expressing the indicated HA-tagged TRIM5 constructs from (E). *n* = 3 biological replicates, each with >100 cells.

**Figure 4.**
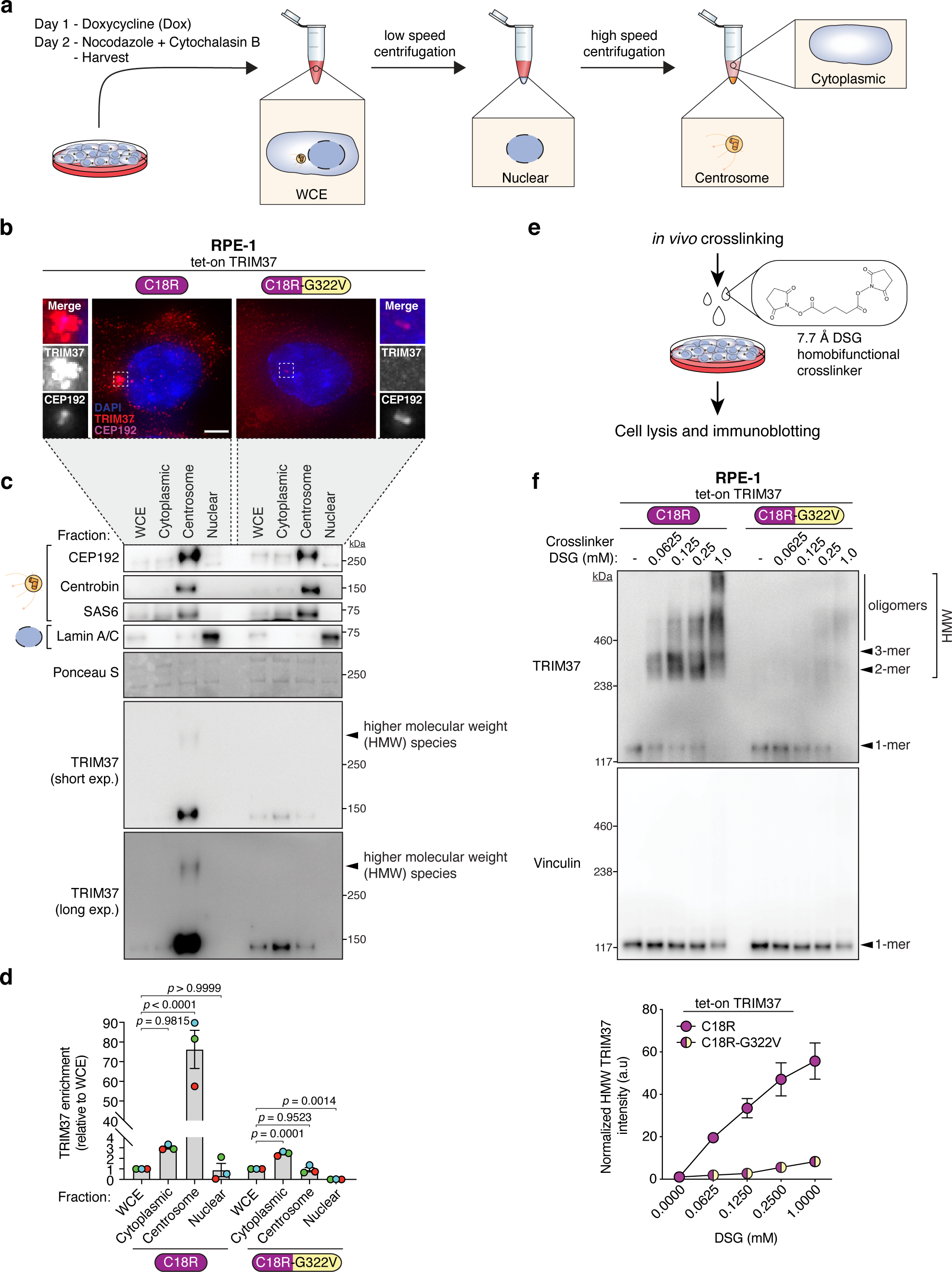
TRAF-directed higher-order assembly of TRIM37 at the centrosome. (A) Experimental schematic of the centrosome-enrichment assay used to separate nuclear, cytoplasmic, and centrosomal fractions, as analysed in (C-D). (B) Representative images of RPE-1 cells expressing the TRIM37 RING domain mutant (C18R) or TRIM37 RING-TRAF double mutant (C18R-G322V). *n* = 3 biological replicates. Scale bars, 5 μm. (C) Immunoblot showing TRIM37 protein levels across the indicated cellular fractions. Validation markers include CEP192, Centrobin, and SAS6 for the centrosomal fraction and Lamin A/C for the nuclear fraction. Ponceau-stained blot indicates loading. Representative data; *n* = 3 biological replicates. WCE, whole-cell extract; exp, exposure. (D) Densitometric analysis of immunoblot in (C) with graph depicting TRIM37 enrichment in indicated fractions relative to WCE. *n* = 3 biological replicates. *P* values, one-way ANOVA with post hoc Dunnett’s multiple comparisons test to evaluate enrichment of TRIM37 in each cellular fraction relative to WCE. Mean ± s.e.m. (E) Schematic of the *in vivo* crosslinking protocol applied to RPE-1 cells using membrane-permeable crosslinkers to elucidate TRIM37 oligomerization dynamics. (F) Top, immunoblot showing detection of various higher molecular weight (HMW) species of TRIM37 upon treatment with increasing concentrations of DSG crosslinker. Vinculin is used as a loading and oligomerization control. Representative data; *n* = 3 biological replicates. Bottom, Densitometric analysis of immunoblot with a graph depicting normalized HMW TRIM37 intensity upon increasing DSG concentrations relative to DMSO control (−DSG). Mean ± s.e.m.

### TRIM37 undergoes centrosome-templated oligomerization

We posited that TRIM37 forms higher-order assemblies crucial for centrosome regulation. Immunoblotting of HA-TRIM37 in RPE-1 whole-cell lysates revealed a band migrating at ∼130 kDa corresponding to monomeric TRIM37, and a distinct higher molecular weight (HMW) species formed by the RING (C18R) TRIM37 mutant that migrated >250kDa (Extended Data Fig. 2a). The TRIM37 HMW species was absent in the MUL B-box (C109S) or TRAF (G322V) mutants (Extended Data Fig. 2a). We deduced that catalytic-dead (C18R) TRIM37 undergoes substrate-templated oligomerization but does not autodegrade, explaining the presence of HMW protein. Consistently, proteasomal inhibition with MG132 prevented the self-degradation of WT TRIM37 and enabled the formation of HMW species (Extended Data Fig. 2b). To investigate if the HMW species of TRIM37 is enriched at the centrosome, we purified centrosomes from RPE-1 cells expressing the RING (C18R) or RING-TRAF (C18R-G322V) double mutant of HA-TRIM37 (Fig. 4a,b). We observed a significant enrichment of monomeric and HMW TRIM37 C18R in the centrosomal fraction compared to the cytoplasmic and nuclear fractions (Fig. 4c,d). Conversely, TRIM37 C18R-G322V did not localize to the centrosome and remained primarily in the cytoplasmic fraction (Fig. 4b–d), suggesting that TRAF-mediated centrosome targeting is required for TRIM37 HMW formation.

We reasoned that the TRIM37 HMW species arises from the incomplete breakdown of the TRIM37 oligomer under denaturing SDS-PAGE conditions. To preserve the substrate-driven assemblies of TRIM37 we conducted *in vivo* crosslinking experiments using two crosslinkers with distinct linker arm lengths (DSG - 7.7 Å and DSS - 11.4 Å, Fig. 4e and Extended Data Fig. 2c). The addition of either crosslinking agent led to a concentration-dependent reduction in the free TRIM37 C18R monomer and corresponding enrichment of TRIM37 HMW forms, including putative dimers, trimers, and beyond—collectively termed oligomers (Fig. 4f and Extended Data Fig. 2c). This effect was not observed with the ubiquitously expressed protein vinculin, ruling out nonspecific crosslinking activity (Fig. 4f and Extended Data Fig. 2c). Notably, the TRIM37 double mutant (C18R-G322V) with a defective TRAF domain did not display clear stabilization of TRIM37 HMW forms with either crosslinking agent (Fig. 4f and Extended Data Fig. 2c).

To extend our findings to endogenous TRIM37, we introduced the TRIM37 (C18R) RING mutation into RPE-1 cells using CRISPR–Cas9 (hereafter referred to as *TRIM37^C18R^*, Extended Data Fig. 2d). Fractionation assays confirmed the centrosomal accumulation of monomeric and HMW forms of endogenous TRIM37^C18R^ (Extended Data Fig. 2e). Moreover, crosslinking-dependent stabilization of additional endogenous HMW species was evident in *TRIM37^C18R^* cells (Extended Data Fig. 2f). Overall, these findings provide support for the hypothesis that substrate binding induces the formation of higher-order TRIM37 assemblies at the centrosome.

### Autodegradation of TRIM37 at the centrosome impedes immunofluorescence detection

Detecting endogenous TRIM37 at the centrosomes has been challenging, with one study revealing no noticeable differences in TRIM37 immunostaining between control and TRIM37 knockout (KO) cells despite using ten different commercial antibodies^18^. Our model posits that TRIM37 undergoes oligomerization at the centrosome, triggering E3 ligase activation followed by autodegradation. Thus, we hypothesized that inactivating the RING domain would reveal stable TRIM37 enrichment at the centrosome. To test this, we assessed TRIM37 protein levels and localization across a panel of cell lines using two commercial antibodies (Extended Data Fig. 3a). Consistent with prior data, we observed a diffuse and punctate pattern of endogenous WT TRIM37 in RPE-1 cells (Extended Data Fig. 3b–g), with weak co-localization at the centrosome (indicated by a yellow arrow in Extended Data Fig. 3c). RING inactivation in RPE-1 *TRIM37^C18R^* cells led to intense TRIM37 centrosome staining (Extended Data Fig. 3b–g). This strong signal was not attributable to increased total protein levels, as 17q23-amplified MCF7 cells overexpress TRIM37 to a higher level than the *TRIM37^C18R^*cells yet lack a comparable centrosomal signal (Extended Data Fig. 3b–g). We conclude that the autodestruction of clustered TRIM37 likely explains the difficulty in detecting the centrosome-localized protein.

### TRIM37 B-box domain mediates higher-order assembly

The B-box domain of TRIM5 is known to drive higher-order assembly via homotrimer formation, with each B-box originating from one TRIM5 dimer (Fig. 5a). These B-box-B-box interactions are mediated in part by the indole sidechains of a key tryptophan residue and are pivotal for facilitating avid binding to retroviral capsids^37,38^ (Fig. 5a). Structural examination of the TRIM37 B-box homotrimer (AlphaFold model) with that of TRIM5 (PDB:5VA4^38^), along with sequence alignment analysis, suggested that the critical tryptophan residue in the TRIM5 B-box corresponds to W120 in TRIM37 (Fig. 5a,b). This analysis also predicted that the MUL C109S patient mutation impacts a Zn-coordinating cysteine residue that is crucial for the proper folding of the B-box domain (Fig. 5b)^39^. Despite similar levels of expression (Fig. 5c), the (C18R-W120E) and (C18R-C109S MUL) B-box mutants of TRIM37 exhibited strongly reduced accumulation at the centrosome and impaired crosslink-stabilized HMW species compared to the RING (C18R) mutant alone (Fig. 5d–f). The addition of nocodazole to induce microtubule (MT) depolymerization before induction of TRIM37 expression led to the formation of multiple discrete cytoplasmic puncta of TRIM37 C18R that did not localize to the centrosome (Fig. 5d).

**Figure 5.**
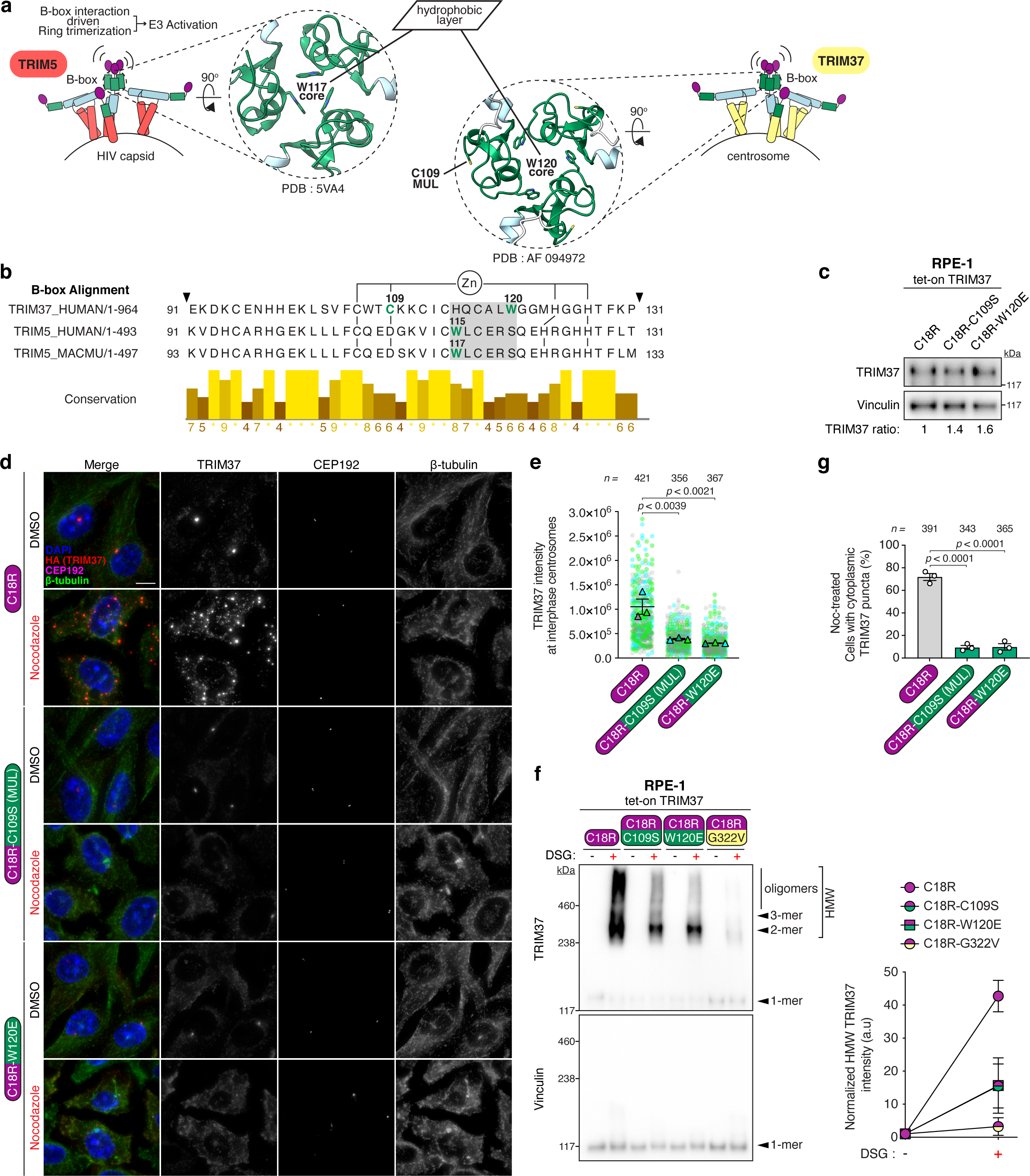
B-box domain mutations impair TRIM37 higher-order assembly. (A) Left, diagram illustrating the B-box trimerization interface of TRIM5 dimers on the HIV capsid. Trimers are stabilized by W117 residues within the hydrophobic core, as shown in the magnified top-down view of the TRIM5 B-box crystal structure (PDB 5VA4). Right, analogous diagram representing a putative oligomer formed by TRIM37 dimers at the centrosome, where B-box domain trimerization is hypothesized to be stabilized by W120 residues, the synonymous counterpart to TRIM5’s W117. A magnified top-down view shows the putative TRIM37 B-box trimer modelled by fitting AlphaFold-predicted TRIM37 monomers onto the TRIM5 crystal structure. (B) Comparative alignment of the B-box domains from human TRIM37 and human and rhesus macaque (*Macaca mulatta*) TRIM5. Residue Cys109, mutated in MUL disease, is pivotal for zinc (Zn) coordination. The highlighted region in grey denotes the sequence alignment where W115/W117 residues in TRIM5 correspond to the W120 residue in TRIM37, signifying a conserved structural motif critical for higher-order assembly. (C) Immunoblot showing total protein expression levels of TRIM37 variants in RPE-1 tet-on TRIM37 cells from (D). Vinculin, loading control. Representative data; *n* = 3 biological replicates. (D) Representative images of RPE-1 tet-on TRIM37 cells expressing the RING domain mutant TRIM37(C18R) or RING-B-box double mutants (C18R-C109S and C18R-W120E). Cells were treated with DMSO (control) or nocodazole (3.3 μM) 30 min before doxycycline induction to depolymerize microtubules. *n* = 3 biological replicates. Scale bars, 5 μm. (E) Quantification of centrosomal TRIM37 signal in DMSO-treated RPE-1 tet-on TRIM37 cells expressing the indicated TRIM37 variants from (D). *n* = 3 biological replicates, each with >100 cells. *P* values, one-way ANOVA with post hoc Dunnett’s multiple comparisons test to evaluate differences between each of the TRIM37 RING-B-box double mutants (C18R-C109S and C18R-W120E) and the RING mutant (C18R). Mean ± s.e.m. (F) Left, immunoblot showing detection of higher molecular weight (HMW) species of indicated TRIM37 variants upon *in vivo* DSG crosslinking. Vinculin is used as a loading and oligomerization control. Representative data; *n* = 3 biological replicates. Right, Densitometric analysis of immunoblot with graph depicting normalized HMW TRIM37 intensity with DSG crosslinker (+) relative to DMSO control (−DSG). Mean ± s.e.m. (G) Evaluation of cytoplasmic TRIM37 puncta prevalence in nocodazole (Noc)-treated RPE-1 tet-on TRIM37 cells expressing the indicated TRIM37 variants from (D). *n* = 3 biological replicates, each with >100 cells. *P* values, one-way ANOVA with post hoc Dunnett’s multiple comparisons test to evaluate differences between each of the TRIM37 RING-B-box double mutants (C18R-C109S and C18R-W120E) and the RING mutant (C18R). Mean ± s.e.m.

These puncta are akin to cytoplasmic bodies (CBs) formed by TRIM5 upon self-association (non-templated assembly)^40^, a process that is also B-box dependent^41,42^. Importantly, the formation of nocodazole-induced TRIM37 cytoplasmic puncta was significantly impaired in cells expressing the two TRIM37 B-box mutants (C18R-C109S and C18R-W120E) (Fig. 5d,g), underscoring the B-box’s crucial role in orchestrating both templated and non-templated, higher-order assembly of TRIM37.

TRIM37 possesses a long, unstructured C-terminal tail following the TRAF domain of unknown function. To evaluate the contribution of this segment (residues 448–964) to TRIM37’s activity and higher-order assembly, we engineered a truncated version of TRIM37 lacking the C-terminus, hereafter referred to as miniTRIM37 (Extended Data Fig. 4a). miniTRIM37 maintained E3 ubiquitin ligase activity, as evidenced by its centrosomal localization and effective degradation of CEP192 (Extended Data Fig. 4b–d). Additionally, miniTRIM37 exhibited oligomerization properties in crosslinking experiments, showing that the core domains (RBCC-TRAF), but not the unstructured C-terminus, are sufficient for TRIM37’s higher-order assembly (Extended Data Fig. 4e).

### Oligomerization is required for the synthetic lethal relationship between TRIM37 overexpression and PLK4 inhibition in 17q23-amplified cancer cells

Prior work has shown that the overexpression of TRIM37 in 17q23-amplified cancers suppresses the formation of PCM foci critical for acentrosomal cell division, thereby explaining the vulnerability of these cancers to centrosome depletion via PLK4 inhibition^24^. As previously reported, treatment of MCF-7 cells with a PLK4 inhibitor resulted in greatly reduced clonogenic viability (Extended Data Fig. 5b), defective spindle assembly, and reduced formation of non-centrosomal PCM foci (Extended Data Fig. 5c,d). These adverse effects were all rescued by TRIM37 knockdown (KD) (Extended Data Fig. 5a–d). To specifically disrupt TRIM37 higher-order assembly, we introduced the C109S B-box mutation into the *TRIM37* gene in MCF-7 cells using CRISPR–Cas9. Although complete allelic conversion of the amplified *TRIM37* gene could not be achieved, sequencing revealed approximately half of MCF-7 *TRIM37* alleles incorporated the C109S variant (hereafter referred to as MCF-7 *TRIM37*^C109S^, Extended Data Fig. 5e).

Importantly, *TRIM37*^C109S^ cells displayed marked resistance to PLK4 inhibition along with a corresponding improvement in the fidelity of mitotic spindle assembly in acentrosomal conditions (Extended Data Fig. 5b–d). This occurred even though *TRIM37*^C109S^ cells expressed similar levels of TRIM37 protein as wild-type MCF-7 cells (Extended Data Fig. 5a), implying that while abundant, TRIM37^C109S^ proteins are functionally defective. These findings highlight the critical role of TRIM37’s oligomerization in driving synthetic lethality with PLK4 inhibition in 17q23 amplified cancers.

### Substrate-induced clustering activates TRIM37

Substrate-induced clustering is a key activation mechanism observed in several members of the TRIM protein family^43^. To directly test the role of clustering in regulating TRIM37 activity, we developed an optogenetic approach to enable spatiotemporal control of TRIM37 clustering independent of its conventional substrate interactions. We fused the TRIM37 TRAF mutant (G322V), impaired in substrate binding, to the fluorescent reporter mNeonGreen (mNG) and CRY2clust, a variant of the Cryptochrome 2 photoreceptor known for its rapid oligomerization upon blue light (BL) exposure^44^ (Extended Data Fig. 6a). Live-cell imaging demonstrated that TRIM37^G322V^-mNG-CRY2 formed cytoplasmic clusters following BL stimulation (Extended Data Fig. 6b,c and Supplementary Video 1). These clusters dissolved over time despite continuous BL exposure, suggesting that clustering triggers TRIM37 autodegradation. Consistently, we observed a marked reduction in whole-cell TRIM37^G322V^-mNG-CRY2 protein levels three hours following BL exposure (Extended Data Fig. 6d).

Proteasome inhibition with MG132 prevented both the time-dependent loss of TRIM37^G322V^-mNG-CRY2 clusters and the decline in protein levels observed following BL stimulation (Extended Data Fig. 6b–d and Supplementary Video 2). Importantly, we also observed the emergence of HMW TRIM37^G322V^-mNG-CRY2 in the presence of MG132 and blue light (Extended Data Fig. 6d, compare Lane 5 vs Lane 4), implying that CRY2-induced clustering triggers TRIM37 oligomerization and subsequent autodegradation via the proteasome pathway.

Having shown that TRIM37 can be activated through clustering independent of substrate binding, we extended our investigation to determine the effect of substrate-driven clustering on TRIM37 activity. We fused mCherry-CRY2 to the C-terminal unstructured region of Centrobin (residues 567–836) (Fig. 6a), identified as a TRIM37-interacting region (data not shown).

**Figure 6.**
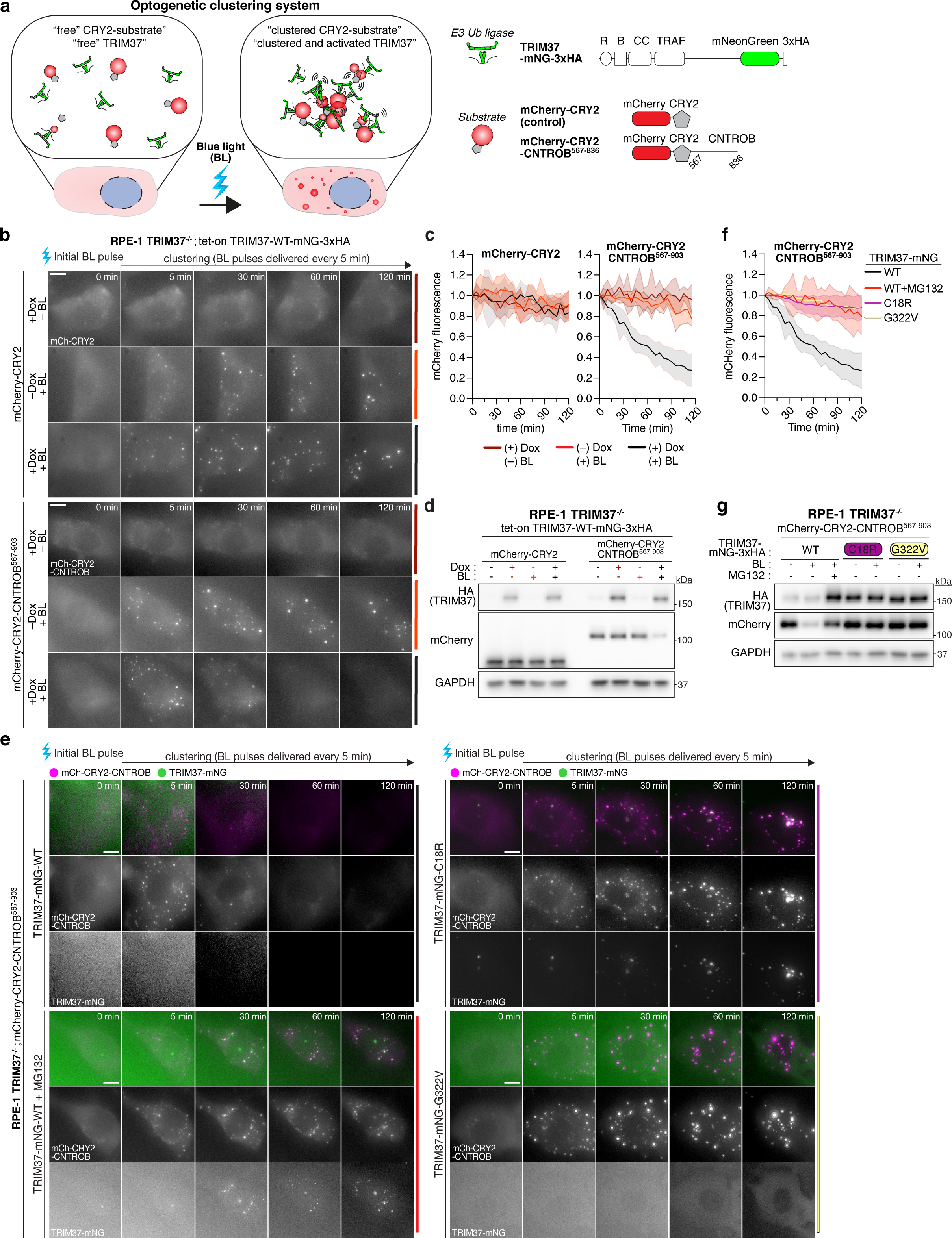
Optogenetic clustering of centrosomal substrates triggers recognition and activation of TRIM37. (A) Left, schematic depicts the blue light (BL)-triggered optogenetic system designed to cluster TRIM37’s cognate centrosomal substrates, enabling the investigation of TRIM37 recognition and activation requirements. Right, schematic of constructs used in the optogenetic experiments, including mNeonGreen-tagged TRIM37 for visualizing recruitment to centrosomal substrates, mCherry-CRY2 fused to Centrobin’s C-terminal unstructured region (residues 567-836) and a mCherry-CRY2 control. (B) Representative time-lapse images of RPE-1 TRIM37^-/-^ cells integrated with optogenetic constructs detailed in (A), incubated with or without doxycycline (Dox), in the absence or presence blue light. Timestamps indicate minutes post blue light exposure. Representative data; *n* = 3 biological replicates. Scale bar = 10 µm. (C) Quantification of mCherry fluorescence intensity from (B), with each condition comprising >30 cells. Mean ± s.d. (D) RPE-1 TRIM37^-/-^ cells integrated with optogenetic constructs detailed in (A) were incubated with or without doxycycline (Dox), in the absence or presence blue light for 3 h prior to immunoblotting for the indicated proteins. GAPDH, loading control. Representative data; *n* = 3 biological replicates. (E) Representative time-lapse images of RPE-1 TRIM37^-/-^ cells integrated with optogenetic constructs and co-expressing different TRIM37 mutants with or without MG132 (10 μM) in the absence or presence blue light. Timestamps indicate minutes post blue light exposure. Representative data; *n* = 3 biological replicates. Scale bar = 10 µm. mCh, mCherry. (F) Quantification of mCherry fluorescence intensity from (E), with each condition comprising >30 cells. Mean ± s.d. (G) RPE-1 TRIM37^-/-^ cells expressing indicated optogenetic constructs and different TRIM37 mutants were treated with or without MG132 (10 μM) in the absence or presence of blue light for 3 h prior to immunoblotting for the indicated proteins. GAPDH, loading control. Representative data; *n* = 3 biological replicates.

TRIM37 displayed no degradation capability towards mCherry-CRY2 alone, either in a diffuse cytosolic state (-BL) or a clustered state (+BL) (Fig. 6b–d and Supplementary Videos 3-5). Illumination with blue light led to the rapid assembly of mCherry-CRY2-Centrobin^567–837^ clusters. Importantly, these clusters dissolved following the induced expression of TRIM37 (Fig. 6b,c and Supplementary Videos 6-8), indicating targeted degradation of the clustered, mCherry-tagged substrate. This was corroborated by a substantial decline in mCherry-CRY2-CNTROB^567– 837^ protein levels only in cells expressing TRIM37 and stimulated with blue light (Fig. 6d). These results demonstrate TRIM37’s degradation activity is specifically directed towards substrates that exist in a clustered configuration. The degradation of mCherry-CRY2-CNTROB^567–837^ clusters by TRIM37 required its E3 ligase function for proteasomal degradation and TRAF domain for substrate recruitment: the C18R RING mutant localized to but failed to degrade the mCherry-CRY2-CNTROB^567–837^ clusters, whereas the G322V TRAF mutant was not recruited to the clusters and could not degrade them (Fig. 6e–g and Supplementary Videos 9-12). These findings support a model where TRIM37 is activated through substrate-induced clustering leading to the degradation of the entire TRIM37-substrate complex.

## Discussion

### Mechanistic dissection of MTOC regulation by TRIM37

In this study, we elucidate the mechanism underlying TRIM37’s activation and its role in the regulation of microtubule-organizing centers (MTOCs). Our findings identify centrosomes, Centrobin assemblies, and non-centrosomal PCM foci as platforms for TRIM37 activation.

Guided by prior work on TRIM5, we demonstrate that TRIM37 E3 ligase activation occurs following TRAF-domain directed clustering of multiple TRIM37 dimers on its substrate (Fig. 7) (i). Upon binding, the oligomerization of TRIM37, mediated by its B-box domain, leads to higher-order structural integrity and increased substrate avidity (ii). The interaction of the B-box domains facilitates the dimerization of RING domains from neighbouring TRIM37 molecules (iii), culminating in E3 ligase activation and the subsequent degradation of both substrate and TRIM37 (iv).

**Figure 7.**
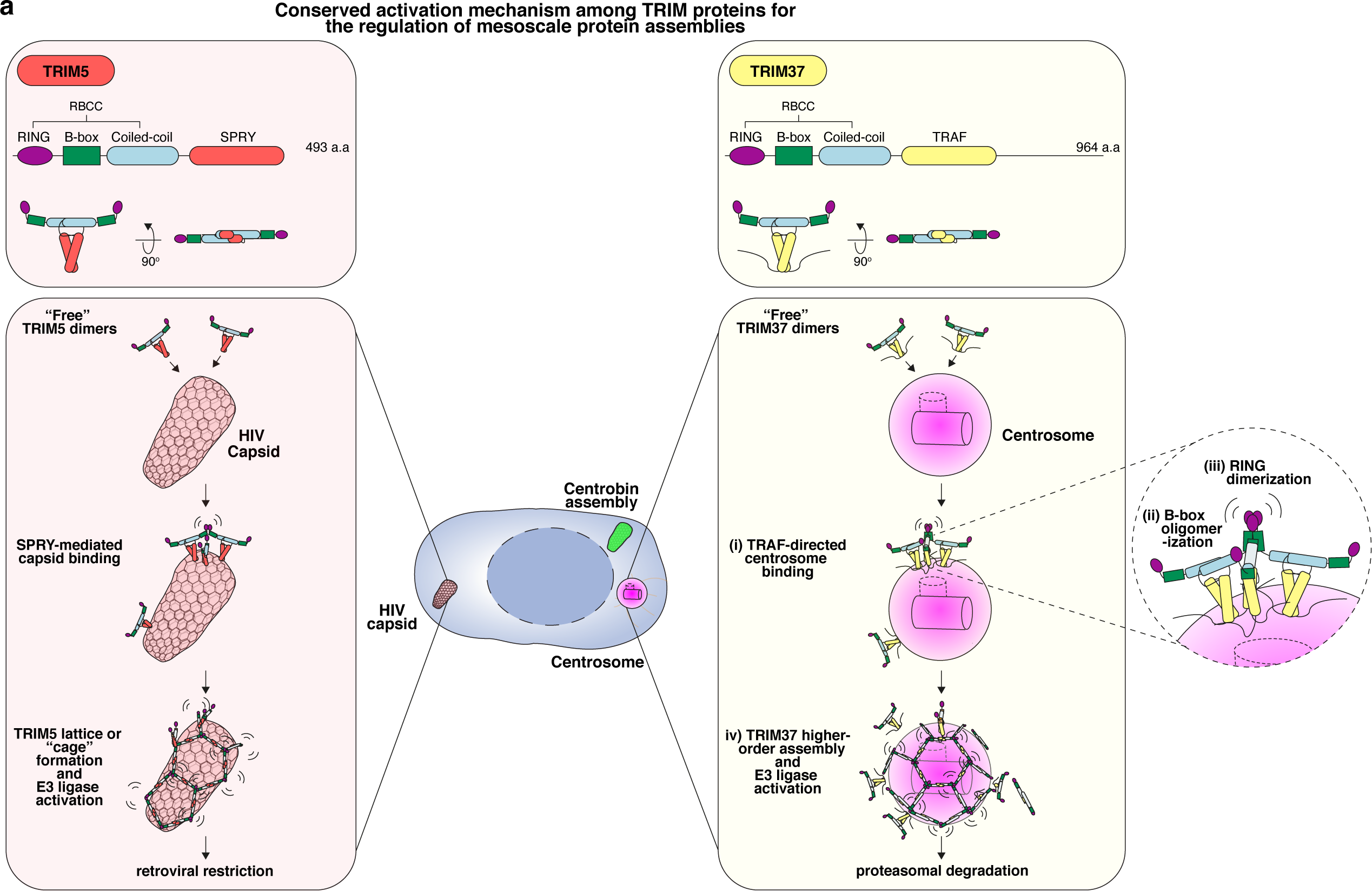
TRIM37 regulates MTOC function via substrate-templated higher-order assembly. (A) Model illustrating how TRIM37 regulates MTOCs through substrate-templated higher-order assembly, demonstrated here using centrosomes, highlighting a conserved mechanism reminiscent of TRIM5’s role in HIV capsid restriction.

The direct coupling of E3 activation to substrate assembly state allows TRIM37 to discriminate between soluble monomeric proteins and those organized into higher-order structures (Fig. 6). This strategy ensures TRIM37’s regulatory activities are confined to functional MTOCs while allowing cells to maintain a pool of centrosomal building blocks. The autodegradation of TRIM37, along with its substrate, limits the enzyme’s degradative capacity and may explain why TRIM37 does not achieve complete degradation of centrosome-associated PCM. Additional mechanisms might shield centrosome-incorporated proteins from TRIM37-mediated turnover, thereby maintaining centrosome homeostasis. The activity of kinases driving PCM expansion or deubiquitinases enlisted by other centrosomal proteins may provide such protection. The suppression of non-centrosomal MTOCs by TRIM37 ensures the centrosome’s exclusive role during cell division, thereby safeguarding mitotic accuracy.

### Pathological and therapeutic insights informed by the TRIM37 activation model

Our model provides a framework for understanding TRIM37’s role in two human conditions associated with centrosome dysfunction. We demonstrate how Mulibrey nanism patient mutations in *TRIM37* compromise its E3 ligase functionality, giving rise to the formation of pathological Centrobin-scaffolded assemblies. Specifically, the G322V mutation in the TRAF domain prevents substrate engagement necessary for TRIM37’s assembly-driven activation, while the C109S mutation in the B-box domain impairs its oligomerization capability.

In cancers characterized by 17q23 amplification, TRIM37 overexpression impedes the assembly of PCM foci that are critical for cells undergoing acentrosomal mitosis^24,25^. This vulnerability underpins an ongoing phase I clinical trial (NCT06232408) utilizing a PLK4 inhibitor to induce centrosome loss for cancer killing. Our optogenetic experiments provide additional insight into this synthetic lethal interaction, revealing that the coalescence of centrosomal proteins triggers TRIM37 activation and rapid degradation of these foci (Fig. 6). Consistently, B-box mutations that impair TRIM37 oligomerization dramatically reduced the sensitivity of 17q23-amplified cancer cells to PLK4 inhibitor treatment (Extended Data Fig. 5). Additionally, we show that TRIM37’s presence at the centrosome is obscured by its autodegradation (Extended Data Fig. 3). This has implications for patient stratification strategies that leverage TRIM37 overexpression as a biomarker to identify tumors susceptible to PLK4 inhibition. Our data indicate that assessing mRNA expression or total protein detection should be prioritized over immunohistochemistry protocols when evaluating TRIM37 expression levels.

### A unifying paradigm of TRIM Proteins in mesoscale assembly regulation

Our work highlights that members of the TRIM protein family with the core RBCC domain architecture deploy an evolutionarily conserved strategy for substrate regulation. The acquisition of the TRAF domain^26^–absent in other TRIM proteins–marks a key event that enabled the recognition and regulation of centrosome substrates in higher-order configurations^45–48^, thus establishing TRIM37 as the principal MTOC regulator within the TRIM superfamily.

While this role diverges from the classical antiviral functions ascribed to TRIM proteins like TRIM5 and TRIM21^29,49^, our findings suggest a unifying paradigm in which TRIM proteins regulate a spectrum of higher-order assemblies, ranging from extrinsic viral entities to intrinsic cellular structures (Fig. 3 and Fig. 7). Echoing its antiviral relatives, we consider TRIM37 as a ‘restriction factor’ for non-centrosomal MTOCs. This concept is complemented by the recently identified role of TRIM11 in mitigating TAU aggregation in Alzheimer’s disease^50^, where we speculate that the assembly of TAU fibrils may act as the trigger for TRIM11-mediated degradation. These insights could lay the groundwork for developing TRIM-based PROTAC strategies that selectively target pathological assemblies while sparing monomeric proteins.

## Acknowledgements

This work was supported by research grants R01GM114119, R01GM133897 and R01CA266199 (to A.J.H) from the National Institutes of Health. We thank Miguel R. Leung and Niladri K. Sinha for providing valuable experimental insights.

## Author contributions

Z.Y.Y. designed, performed, and analysed the majority of the experiments and prepared the figures. S.S. assisted with cloning and immunofluorescence analysis. M.v.B performed experiments to identify TRIM37 binding regions within Centrobin. Z.Y.Y. and A.J.H. conceived the study. A.J.H. supervised the study. Z.Y.Y. and A.J.H. co-wrote the manuscript.

## Declaration of interests

The authors declare no competing interests.

**Correspondence and requests for materials** should be addressed to Z.Y.Y and A.J.H.

## Supplementary video legends

**Supplementary Video 1. (related to Extended Data Figure 6)**

Time-lapse of an RPE-1 cell expressing the optogenetic TRIM37G322V-mNeonGreen-CRY2 construct, with blue light (470 nm filter) pulses applied during imaging intervals to induce and sustain CRY2 clustering and to visualize TRIM37-mNeonGreen dynamics. Timestamps indicate minutes from the initial blue light exposure.

**Supplementary Video 2. (related to Extended Data Figure 6)**

Time-lapse of an RPE-1 cell expressing the optogenetic TRIM37G322V-mNeonGreen-CRY2 construct, with blue light (470 nm filter) pulses applied during imaging intervals to induce and sustain CRY2 clustering and to visualize TRIM37-mNeonGreen dynamics. Cells were incubated with MG132 1 hour prior to blue light exposure. Timestamps indicate minutes from the initial blue light exposure.

**Supplementary Video 3. (related to Figure 6)**

Time-lapse of an RPE-1 TRIM37^-/-^ cell expressing the optogenetic mCherry-CRY2 construct, incubated with doxycycline (+Dox), but in the absence blue light (–BL). Timestamps indicate minutes from the first imaged frame. mCherry-CRY2 is displayed in grayscale.

**Supplementary Video 4. (related to Figure 6)**

Time-lapse of an RPE-1 TRIM37^-/-^ cell expressing the optogenetic mCherry-CRY2 construct, subjected to blue light pulses (+BL) at each imaging interval, but incubated without doxycycline (–Dox). Timestamps indicate minutes from the initial blue light exposure. mCherry-CRY2 is displayed in grayscale.

**Supplementary Video 5. (related to Figure 6)**

Time-lapse of an RPE-1 TRIM37^-/-^ cell expressing the optogenetic mCherry-CRY2 construct, incubated with doxycycline (+Dox) and subjected to blue light pulses (+BL) at each imaging interval. Timestamps indicate minutes from the initial blue light exposure. mCherry-CRY2 is displayed in grayscale.

**Supplementary Video 6. (related to Figure 6)**

Time-lapse of an RPE-1 TRIM37^-/-^ cell expressing the optogenetic mCherry-CRY2-CNTROB^567– 837^ construct, incubated with doxycycline (+Dox), but in the absence blue light (–BL). Timestamps indicate minutes from the first imaged frame. mCherry-CRY2-CNTROB^567–837^ is displayed in grayscale.

**Supplementary Video 7. (related to Figure 6)**

Time-lapse of an RPE-1 TRIM37^-/-^ cell expressing the optogenetic mCherry-CRY2-CNTROB^567– 837^ construct, subjected to blue light pulses (+BL) at each imaging interval, but incubated without doxycycline (–Dox). Timestamps indicate minutes after initial blue light exposure. mCherry-CRY2-CNTROB^567–837^ is displayed in grayscale.

**Supplementary Video 8. (related to Figure 6)**

Time-lapse of an RPE-1 TRIM37^-/-^ cell expressing the optogenetic mCherry-CRY2-CNTROB^567– 837^ construct, incubated with doxycycline (+Dox) and subjected to blue light pulses (+BL) at each imaging interval. Timestamps indicate minutes from the initial blue light exposure. mCherry-CRY2-CNTROB^567–837^ is displayed in grayscale.

**Supplementary Video 9. (related to Figure 6)**

Time-lapse of an RPE-1 TRIM37^-/-^ cell expressing the optogenetic mCherry-CRY2-CNTROB^567– 837^ construct and TRIM37-mNG-WT, subjected to blue light pulses (+BL) at each imaging interval. Timestamps indicate minutes from the initial blue light exposure. mCherry-CRY2-CNTROB^567– 837^ is displayed in magenta, and TRIM37-mNG in green.

**Supplementary Video 10. (related to Figure 6)**

Time-lapse of an RPE-1 TRIM37^-/-^ cell expressing the optogenetic mCherry-CRY2-CNTROB^567– 837^ construct and TRIM37-mNG-WT, subjected to blue light pulses (+BL) at each imaging interval. Cells were incubated with MG132 1 hour prior to blue light exposure. Timestamps indicate minutes from the initial blue light exposure. mCherry-CRY2-CNTROB^567–837^ is displayed in magenta, and TRIM37-mNG in green.

**Supplementary Video 11. (related to Figure 6)**

Time-lapse of an RPE-1 TRIM37^-/-^ cell expressing the optogenetic mCherry-CRY2-CNTROB^567– 837^ construct and TRIM37-mNG-C18R, subjected to blue light pulses (+BL) at each imaging interval. Timestamps indicate minutes from the initial blue light exposure. mCherry-CRY2-CNTROB^567–837^ is displayed in magenta, and TRIM37-mNG in green.

**Supplementary Video 12. (related to Figure 6)**

Time-lapse of an RPE-1 TRIM37^-/-^ cell expressing the optogenetic mCherry-CRY2-CNTROB^567– 837^ construct and TRIM37-mNG-G322V, subjected to blue light pulses (+BL) at each imaging interval. Timestamps indicate minutes from the initial blue light exposure. mCherry-CRY2-CNTROB^567–837^ is displayed in magenta, and TRIM37-mNG in green.

## Methods

### Cell lines and culture conditions

hTERT RPE-1 and MCF-7 cells were grown in DMEM medium (Corning Cellgro) containing 10% fetal bovine serum (Sigma), 100 U/ml penicillin, 100 U/ml streptomycin and 2 mM L-glutamine. All cell lines were maintained at 37°C in a 5% CO_2_ atmosphere with 21% oxygen and routinely checked for mycoplasma contamination.

### Gene targeting and stable cell lines

To generate CRISPR/Cas9-mediated knockout lines, the sgRNA targeting *TRIM37* (*TRIM37Δ*, 5′-ctccccaaagtgcacactga-3′) was cloned into the PX459 vector (#62988; Addgene) containing a puromycin resistance cassette. Cells were transiently transfected (Lipofectamine LTX, Thermo Fisher Scientific) with the PX459 plasmid and positive selection of transfected cells was performed 2 days after transfection with 2.0 ug/ml puromycin. Monoclonal cell lines were isolated by limiting dilution. The presence of gene-disrupting insertions or deletions (indels) in edited cell lines was confirmed via Sanger sequencing, analysed by Tracking of Indels by Decomposition (TIDE: https://tide.nki.nl/)^51^, and the ablation of protein production was assessed by immunoblotting.

To generate TRIM37-overexpressing RPE-1 cell lines, TRIM37 open reading frame (ORF) was cloned into a tet-inducible lentiviral vector containing a C-terminal 3xHA tag. The C18R, C109S and G322V mutations were introduced using PCR-directed mutagenesis and subsequently verified by Sanger sequencing. TRIM37 ΔB-box (residues 91–131 deleted), ΔTRAF (residues 274–448 deleted), and miniTRIM37 (residues 459–964 deleted) were constructed using Gibson assembly and verified by Sanger sequencing. Lentiviral particles were produced as described below. Cells were transduced and stable polyclonal populations of cells selected and maintained in the presence of 1.0 µg/mL puromycin.

To generate TRIM5-WT or TRIM5 chimera expressing RPE-1 cell lines, the TRIM5 ORF (#79066; Addgene) was PCR amplified and cloned into a tet-inducible lentiviral vector containing a C-terminal 3xHA tag. The TRIM5-TRAF chimera was engineered by replacing the SPRY domain (residues 303–493) with TRIM37’s TRAF domain (residues 274–448) using Gibson assembly, with the constructs verified by Sanger sequencing. Lentiviral particles were produced as described below. Cells were transduced and stable polyclonal populations of cells selected and maintained in the presence of 1.0 µg/mL puromycin.

To generate cell lines expressing mCherry-CRY2 variants, the sequence encoding mCherry-CRY2clust (#105624; Addgene) was PCR-amplified. This construct, either fused with the C-terminal region of CNTROB (residues 567–836) or alone, was then incorporated via Gibson assembly into a constitutive lentiviral vector that included blasticidin resistance. Lentiviral particles were produced as described below. RPE-1 TRIM37^-/-^ cells engineered with tet-inducible TRIM37-mNeongGreen were transduced and stable polyclonal populations were selected and maintained in the presence of 30.0 µg/mL blasticidin.

To generate RPE-1 and MCF-7 cell lines with targeted edits to the endogenous TRIM37 loci (C18R and C109S, respectively), a CRISPR–Cas9 knock-in (KI) strategy was employed as previously described^52^. Specifically, Alt-R™ crRNAs targeting *TRIM37* (C18R, 5′-ucauuugu auggagaaauugguuuuagagcuaugcu-3’; and C109S 5′-cuccccaaagugcacacugaguuuuagagcuaugcu-3’, both from IDT) were annealed with tracrRNA (IDT) and subsequently combined with recombinant Alt-R™ S.p. Cas9 Nuclease V3 (IDT). The assembled ribonucleoprotein (RNP) complexes and corresponding single-stranded DNA homology templates (C18R 5′-cttgccttttactcttgattcagtagcctaaactggtggaccttacatcttttactgttttcagagcattgctgaggttttccgatgtttcatccgtatggagaaattgcgcgatgcacgcctgtgtcctcattgctccaaactgtgttg-3’; and for C109S 5′-tccaatttaatttataacttcattttcttttcataatgtatagatgtgaaaatcaccatgaaaaacttagtgtattttgctggacttctaagaagtgtatctgccaccaatgtgcactttggggaggaatggtgagcagaacaaattcag-3’, both from IDT) were nucleofected into cells using the 4D-Nucleofector^TM^ X Unit (Lonza) following the prescribed protocols: RPE-1, EA-104 program, P3 Buffer; MCF-7, EN-130 program, SE Buffer. Post-electroporation, cells were treated with 1 μM NU7441 (Selleck Chemicals) for 48 h to enhance homology-directed repair (HDR) efficacy. Monoclonal cell lines were isolated by limiting dilution, with the specific gene edits confirmed via Sanger sequencing.

### RNA interference

shRNAs targeting TRIM37 (TRIM37-1, 5’-tcgagaatatgatgctgtg-3’) were cloned into the pGIPz (Thermo Fisher Scientific) vector. Stable shRNA-mediated knockdown (KD) cell lines were generated by lentivirus-mediated transduction. Polyclonal populations of cells were subsequently selected and maintained in the presence of puromycin (1.0 µg/mL). Knock down efficiency was assessed by immunoblotting.

### Lentiviral production and transduction

Lentiviral expression vectors were co-transfected into 293FT cells with the lentiviral packaging plasmids psPAX2 and pMD2.G (Addgene #12260 and #12259). Briefly, 3 × 10^6^ 293FT cells were seeded into a Poly-L-Lysine coated 10 cm culture dish the day before transfection. For each 10 cm dish the following DNA were diluted in 0.6 mL of OptiMEM (Thermo Fisher Scientific): 4.5 µg of lentiviral vector, 6 µg of psPAX2 and 1.5 µg of pMD2.G. Separately, 72 µl of 1 µg/µl 25 kDa polyethyleneimine (PEI; Sigma) was diluted into 1.2 mL of OptiMEM, briefly vortexed, and incubated at room temperature for 5 min. After incubation, the DNA and PEI mixtures were combined, briefly vortexed, and incubated at room temperature for 20 min. During this incubation, the culture media was replaced with 17 mL of pre-warmed DMEM + 1% FBS. The transfection mixture was then added drop-wise to the 10 cm dish. Viral particles were harvested 48 h after the media change and filtered through a 0.45 µm PVDF syringe filter. The filtered supernatant was either concentrated in 100 kDa Amicon Ultra Centrifugal Filter Units (Millipore) or used directly to infect cells. Aliquots were snap-frozen and stored at −80°C. For transduction, lentiviral particles were diluted in complete growth media supplemented with 10 µg/mL polybrene (Sigma) and added to cells.

### Chemical inhibitors

MG132 (Sigma) was dissolved in dimethyl sulfoxide (DMSO) and used at a final concentration of 10 μM. Nocodazole (Selleck Chemicals) was dissolved in DMSO and used at a final concentration of 3.3 μM. Centrinone (Tocris Bioscience) was dissolved in DMSO and used at a final concentration of 250 nM.

### Structural modelling and sequence alignment

The structure of monomeric TRIM37 (UniProt ID O94762) was obtained from the AlphaFold Protein Structure Database ^53^. Dimerization of TRIM37 (residues 1–448) was modelled using AlphaFold-Multimer on ColabFold (version 1.5.5)^54,55^ with default settings. Of the five models generated, the one with the highest AlphaFold predicted template modelling score (pTM-score) was selected for this study. Structural visualizations were created with UCSF ChimeraX^56^.

Alignment of the B-box domains of TRIM5 (Rhesus macaque; UniProt ID Q0PF16, human; UniProt ID Q9C035) and TRIM37 (human; UniProt ID O94762) was conducted using Jalview^57^.

### BioID sample preparation, mass spectrometry (MS), and data analysis

To generate cell lines for BioID, puro-sensitive RPE-1 cells were transduced with lentivirus containing tet-inducible miniTurbo control, or various miniTurbo-TRIM37 constructs. After 48 h, cells were selected in 2.0 µg/ml puromycin for 2 d followed by expansion into two 15 cm dishes. Six hours prior to biotin labelling, 1 µg/mL doxycycline was added to induce expression of miniTurbo constructs. The culture medium was then supplemented with 250 µM D-biotin (P212121; prepared as 250 mM stock in DMSO) to initiate labelling of proximity interactors. Samples were collected after 4 h of biotin labelling, transferred to 15 mL conical tubes, and rinsed four times with ice-cold PBS to eliminate excess biotin. Cell pellets were lysed in ∼1.5 mL lysis buffer (all buffer recipes have previously been published^58^) by gentle pipetting followed by sonication. Lysates were then clarified by centrifugation at 16,000 × *g* (15 min, 4 °C). Biotinylated proteins were enriched by incubating 50 µL of streptavidin agarose bead resin (Pierce) with the lysates, rotating overnight at 4°C. Beads were then washed for 10 min each with a series of four wash buffers of decreasing detergent concentrations, followed by two final washes in 50 mM ammonium bicarbonate, and then resuspended in ∼60 µL of the same buffer before freezing for mass spectrometry.

For mass spectrometry preparation, proteins were reduced with 1.75 µL 15 mg/mL DTT in 10 mM TEAB, shaking at 56°C for 50 min. Samples were then cooled to room temperature, the pH adjusted to 8 with 500 mM TEAB buffer, and alkylated with 1.8 µL 36 mg/mL iodoacetamide in 100 mM TEAB for 20 min in the dark. Proteolysis was performed by adding 20 ng/µL trypsin (Promega) and incubating at 37°C overnight. Supernatants were collected, beads washed with 0.1x TFA three times, with washes added to supernatant. The pH was adjusted to acidic range, and peptides desalted on u-HLB Oasis plates, eluted with 60% acetonitrile/0.1% TFA, and dried. A 10% aliquot of desalted peptides was analysed on Nano LC-MS/MS on Q Exactive Plus (Thermo Fisher Scientific) in FTFT mode. MS/MS data were processed with Mascot via PD2.2 against RefSeq2017_83 human species database and a small enzyme and standard (BSA) containing database using the FilesRC option, with a mass tolerance of 3 ppm on precursors and 0.01 Da on fragments, and annotating variable modifications such as oxidation on M, carbamidomethyl C, deamidation NQ, with and without Biotin K. The resulting Mascot.dat files were 1) compiled in Scaffold and 2) processed in PD2.2 to identify peptides and proteins using Percolator as a PSM validator.

Protein hits identified exclusively in miniTurbo-TRIM37 BioID and those whose spectral counts in miniTurbo-TRIM37 (WT and C18R) BioID were at least 2-fold greater than those of mTurbo alone (control) were considered as candidates for TRIM37 interaction. A second criterion was applied whereby hits whose spectral counts in miniTurbo-TRIM37 (WT and C18R) BioID were 2-fold greater than those of miniTurbo-TRIM37 (G322V) were identified as TRAF-specific interactors. Additionally, the minimum spectral count for inclusion was set to two, and common contaminants listed on the CRAPome database^59^ were excluded. The filtered list of BioID hits was annotated with Gene Ontology (GO) terms via the Panther classification system^60^ and analysed using the statistical overrepresentation test (binomial) to derive *P* values^61^. Visualization of data was done using the dot plot generator from ProHits-viz^62^.

### Antibody techniques

For immunoblot analyses, protein samples were resolved by SDS-PAGE on pre-cast NuPAGE™ gels (1.0 mm 4–12% Bis-Tris or 1.5 mm 3–8% Tris-Acetate for HMW TRIM37, Invitrogen) with molecular weight ladders (PageRuler Plus or HiMark pre-stained protein standard for HMW TRIM37, Invitrogen). Following electrophoresis, proteins were transferred onto nitrocellulose membranes using a Mini Trans-Blot Cell (BioRad) wet transfer system and subsequently probed with the following primary antibodies: TRIM37 (rabbit, Bethyl, A301-174A, 1:1000), HA (rat; Roche, ROAHAHA; 1:1000), β-actin (mouse, Santa Cruz Biotechnology, sc-4778, 1:1000), CEP192 (rabbit, raised against CEP192 residues 1–211, home-made^24^, 1:1000), CNTROB (rabbit, Atlas Antibodies, HPA023319, 1:1000), SAS-6 (mouse, Santa Cruz Biotechnology, sc-81431, 1:1000), vinculin (mouse, Santa Cruz Biotechnology, sc-73614, 1:1000), GAPDH (mouse, Santa Cruz Biotechnology, sc-47724, 1:1000), TRIM37 (rabbit, Cell Signaling, #96167, 1:1000, see Extended Data Fig. 3e–g), and mCherry (rabbit, Abcam, ab167453, 1:4000). Detection was performed using HRP-conjugated secondary antibodies: anti-mouse (horse; Cell Signaling, #7076; 1:1000), anti-rat (goat; Cell Signaling, #7077; 1:1000), anti-rabbit (goat; Cell Signaling, #7074, 1:1000), and streptavidin (Cell Signaling, #3999; 1:1500), with SuperSignal West Pico PLUS or Femto Maximum chemiluminescence substrate (Thermo Fisher Scientific). Signals were visualized and acquired using a Genesys G:Box Chemi-XX6 system (Syngene).

For immunofluorescence, cells were cultured on 12-mm glass coverslips and fixed for 8 min in 100% ice-cold methanol at −20°C. Cells were blocked in 2.5% FBS, 200 mM glycine, and 0.1% Triton X-100 in PBS for 1 h. Antibody incubations were conducted in the blocking solution for 1 h. DNA was stained with DAPI, and cells were mounted in ProLong Gold Antifade (Invitrogen). Staining was performed with the following primary antibodies: HA (rat; Roche, ROAHAHA; 1:500), CEP192-Cy5 (directly-labelled goat, raised against CEP192 residues 1–211, home-made, 1:1000), CNTROB (rabbit, Atlas Antibodies, HPA023319, 1:1000), Streptavidin Alexa Fluor™ 555 Conjugate (Invitrogen, S32355, 1:1000), β-tubulin (guinea pig, ABCD Antibodies, ABCD_AA344, 1:4000), TRIM37 (rabbit, Cell Signaling, #96167, 1:1000, see Extended Data Fig. 3e–g), Centrin (mouse, Millipore, 04-1624, 1:1000), and α-tubulin (rat, Invitrogen, MA1-80017, 1:1000).

Immunofluorescence images were acquired using a DeltaVision Elite system (GE Healthcare) controlling a Scientific CMOS camera (pco.edge 5.5). Acquisition parameters were controlled by SoftWoRx suite (GE Healthcare). Images were collected at room temperature (25°C) using an Olympus 40x 1.35 NA, 60x 1.42 NA or Olympus 100x 1.4 NA oil objective at 0.2 μm z-sections. Images were acquired using Applied Precision immersion oil (N=1.516). For quantitation of signal intensity at the centrosome, deconvolved 2D maximum intensity projections were saved as 16-bit TIFF images. Signal intensity was determined using ImageJ by drawing a circular region of interest (ROI) around the centriole (ROI S). A larger concentric circle (ROI L) was drawn around ROI S. ROI S and L were applied to the channel of interest and the signal in ROI S was calculated using the formula IS − [(IL − IS/AL − AS) × AS], where A is area and I is integrated pixel intensity.

### Centrosome enrichment assays

Centrosome purification was performed as described previously^63^, with some modifications. RPE-1 cells seeded at a density of 2 × 10^6^ cells in 15 cm dishes were treated with 1 μg/mL doxycycline (Thermo Fisher Scientific) for 18 h to induce TRIM37 protein expression. Prior to harvest, cells were treated with 3.3 μM nocodazole (Selleck Chemicals) and 5 μg/mL cytochalasin B (Cayman Chemical) for 1 h 15 min to depolymerize microtubule and actin networks, thus facilitating the dissociation of centrosomes from the nuclei. Cells were then washed sequentially with ice-cold 1 × PBS, 8% sucrose in 0.1 × PBS, 8% sucrose in deionized H_2_O, and Tris-HCl (pH 8.0) containing 0.46 μL/mL β-Mercaptoethanol (β-ME) (Sigma). Lysis was carried out at 4 °C with a 1 mM Tris-HCl buffer (pH 8.0) that included 0.5% IGEPAL CA-630 (Sigma), 0.5 mM MgCl_2_, 0.1 mM PMSF (Sigma), 0.1 mM Ortho-vanadate (Sigma), protease and phosphatase cocktail inhibitors (Roche), 0.46 μL/mL β-ME and 10 U/L Benzonase (MilliporeSigma), with vigorous rocking for 15 min. The whole cell lysate was initially centrifuged at 2,500 × *g* (5 min, 4 °C) to isolate the nuclear fraction (pellet). The supernatant was then subjected to ultracentrifugation at 21,100 × g (15 min, 4 °C) to further separate the centrosome-containing fraction (pellet) from the cytosolic fraction.

### *In vivo* crosslinking assays

RPE-1 cells seeded at a density of 9 × 10^5^ cells/well in 12-well plates were treated with 1 μg/mL doxycycline (Thermo Fisher Scientific) for 16 h to induce TRIM37 protein expression. Crosslinking agents DSS (disuccinimidyl suberate) and DSG (disuccinimidyl glutarate) (Thermo Fisher Scientific) were then prepared as solutions in DMSO and added to the culture medium. After a 12-min incubation at room temperature to facilitate crosslinking, the medium containing crosslinkers was removed, and the reaction was quenched by adding a 50 mM Tris-HCl solution (pH 8.0) directly to the wells for an additional 10 min at room temperature. Cell lysates were subsequently harvested, clarified by brief centrifugation at 8000 x *g* (5 min, 4 °C), and prepared for immunoblot analysis.

### Blue light (BL)-induced CRY2 clustering experiments

Fluorescent RPE-1 cell lines were seeded into µ-Slide 4-well or 8-well glass bottom chamber slides (Ibidi). Cells were treated with 1 μg/mL doxycycline, with or without MG132, to induce TRIM37 protein expression one hour before blue light (BL) exposure and were kept in the dark.

Time-lapse imaging was performed using a Zeiss Axio Observer 7 inverted microscope equipped with Slidebook 2023 software (3i—Intelligent, Imaging Innovations, Inc.), X-Cite NOVEM-L LED laser and filter cubes, and a Prime 95B CMOS camera (Teledyne Photometrics) with a 40×/1.3 plan-apochromat oil immersion objective. During imaging, cell conditions were maintained at 37°C, with 5% CO_2_, and 60% relative humidity (RH) using a stage top incubator (Okolab). The 470-nm filter was employed to induce CRY2 clustering and simultaneously image TRIM37-mNeonGreen, while the 555-nm filter was used for mCherry-CRY2 visualization. Images were captured every 5 min in 14 × 2 μm z-sections, and integrated fluorescence intensity measurements (regions of interest were manually delineated to encompass the full area of each individual cell where clustering occurred) were derived from maximum intensity projected 2D time-lapse images in Fiji. Following background subtraction, fluorescence intensity was normalized to the initial image frame (*t* = 0, prior to BL illumination).

For immunoblot analysis, cells seeded into µ-Slide 4-well or 8-well glass bottom chamber slides (Ibidi) were exposed to BL using a DR89X blue LED transilluminator (Clare Chemical) controlled by a programmable timed power switch. The exposure regimen involved cycles of 5 s of BL exposure “on” followed by 5 min “off”, continuing over a total duration of 3 h before the cells were harvested.

### PLK4i survival assays

MCF-7 cells seeded in triplicate at a density of 1.25 × 10^4^ cells/well in 6-well plates were treated with either DMSO (control) or PLK4i (250 nM centrinone) 16 h later. After the indicated number of days, cells were fixed and stained using 0.5% (w/v) crystal violet in 20% (v/v) methanol for 5 minutes. Excess crystal violet was thoroughly rinsed away with distilled water and plates dried overnight. For quantification, bound crystal violet was dissolved in 10% (v/v) acetic acid in dH_2_O and absorbance of 1:50 dilutions were measured at 595 nm using a Synergy HT Microplate Reader (BioTek Instruments Inc). The optical density at 595 nm (OD595) served as a quantitative metric of relative cell growth.

## Data availability

Data that support the findings of this study are available from the corresponding authors upon reasonable request. Source data are provided with the paper.

## Extended Data Figure Legends

**Extended Data Figure 1.**
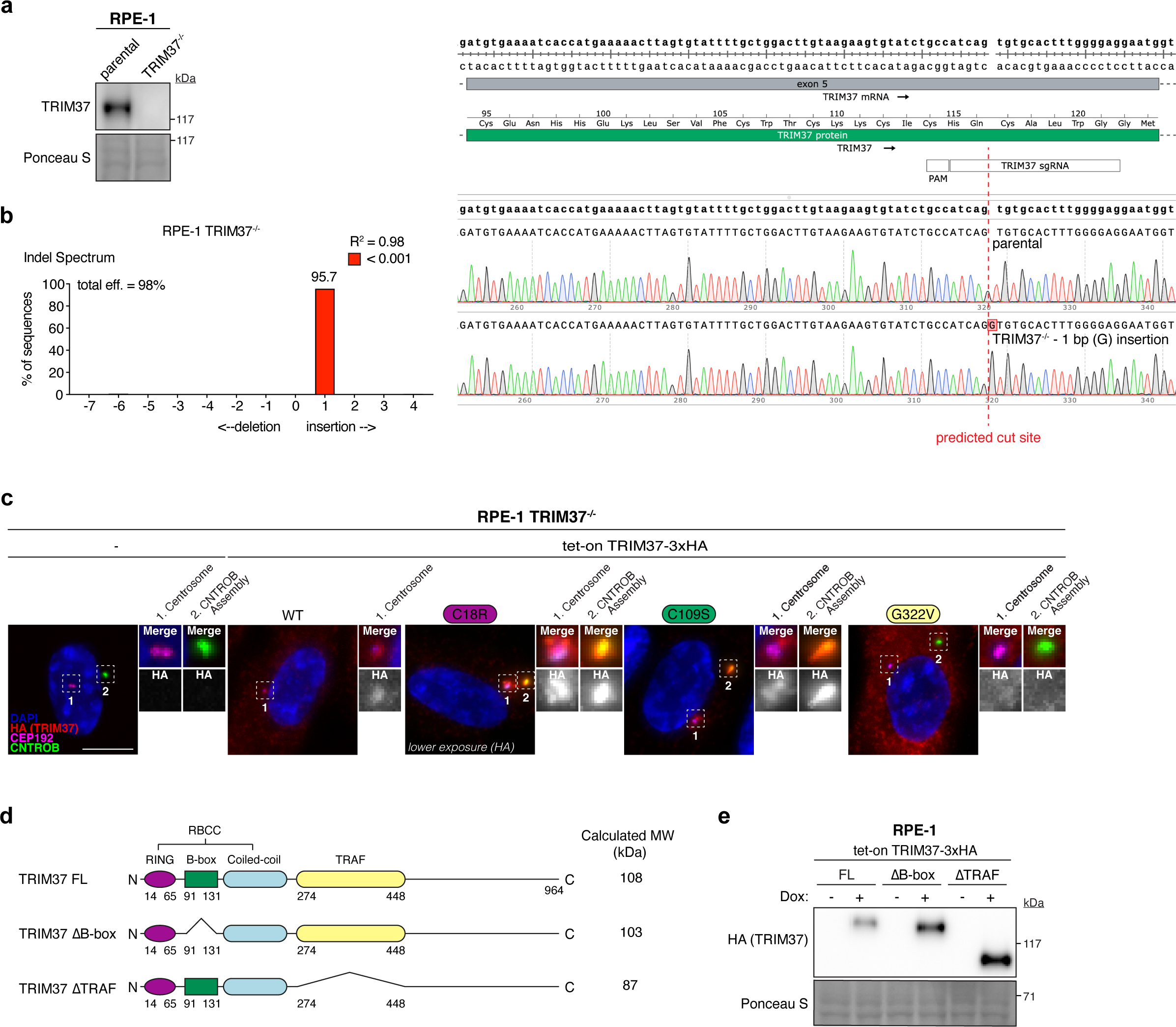
Effect of TRIM37 mutations on the regulation of the centrosome and Centrobin assemblies. (related to Figure 1). (A) Immunoblot showing TRIM37 protein levels in parental and CRISPR–Cas9 edited *TRIM37^−/−^* RPE-1 cells. Ponceau-stained blot indicates loading. Representative data; *n* = 3 biological replicates. (B) Left, Tracking of Indels by Decomposition (TIDE) analysis histogram reveals a one base pair insertion (+1 bp) adjacent to the predicted cut site in the *TRIM37*^−/−^ RPE-1 cell line. Right, representative Sanger sequencing traces used for TIDE analysis, highlighting the +1 bp insertion. (C) Representative images of RPE-1 *TRIM37*^−/−^ cells and those expressing the indicated HA-tagged TRIM37 variants. Inset #1 denotes the centrosome, marked by CEP192, and inset #2 denotes the Centrobin assembly, identified by intense Centrobin staining that is non-centrosome localized. Representative data; *n* = 3 biological replicates. Scale bars, 5 μm. (D) Schematic representation of TRIM37 HA-tagged domain-specific deletion constructs compared to full-length (FL) protein. (E) Immunoblot showing expression levels of FL TRIM37 and the respective deletion mutants in RPE-1 tet-on TRIM37 cells. Ponceau-stained blot indicates loading. Representative data; *n* = 3 biological replicates.

**Extended Data Figure 2.**
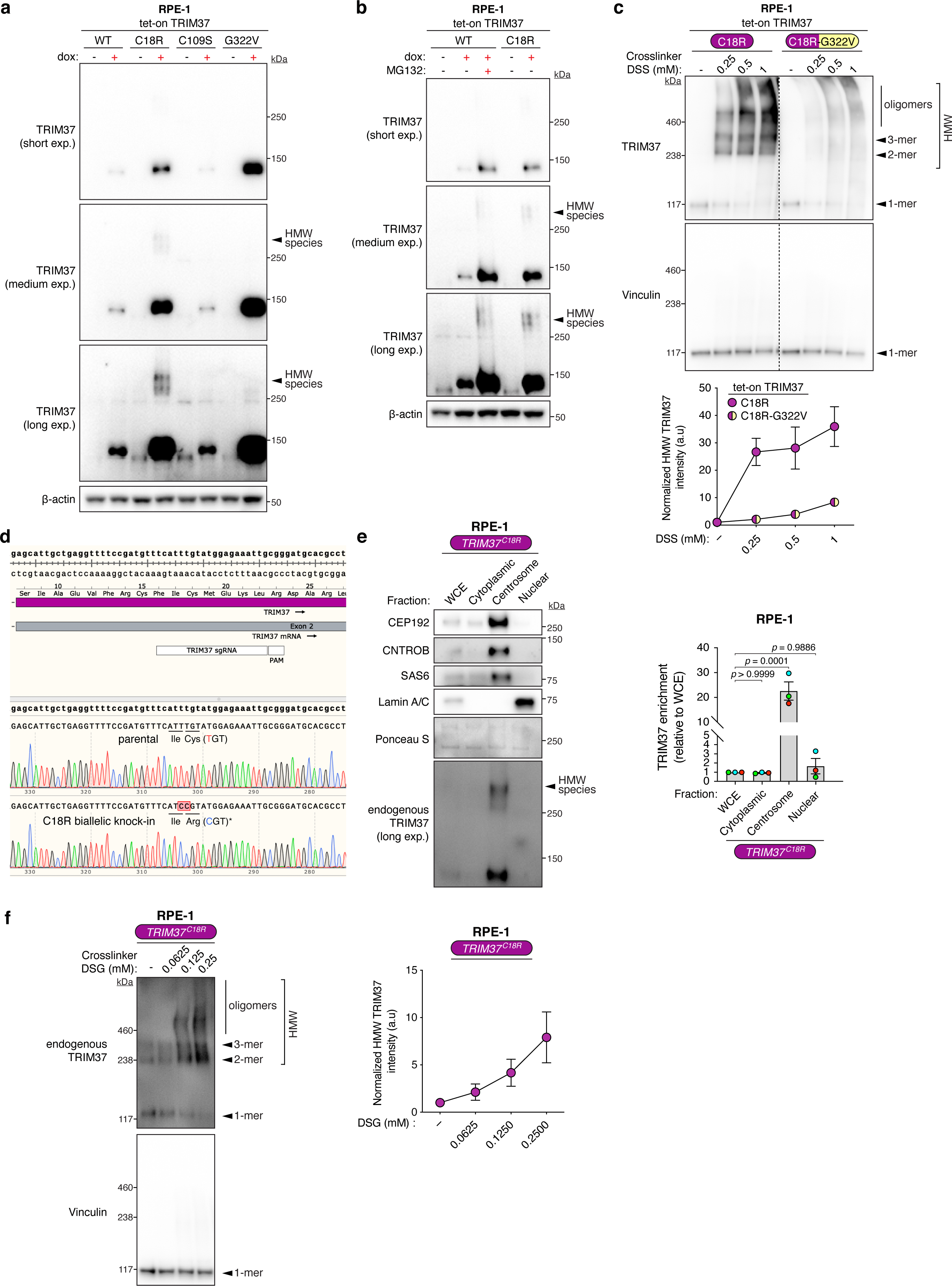
Characterization of higher molecular weight (HMW) TRIM37 species. (related to Figure 4). (A) Immunoblot showing expression levels of wild-type (WT) TRIM37 and indicated mutants in RPE-1 tet-on TRIM37 cells. Higher molecular weight (HMW) TRIM37 species are prominently formed in the C18R mutant and indicated with an arrow. β-Actin, loading control. Representative data; *n* = 3 biological replicates. (B) Same as in (A) but with MG132 (10 μM) treatment to achieve proteasomal inhibition and stabilization of WT TRIM37. β-Actin, loading control. Representative data; *n* = 3 biological replicates. (C) Top, immunoblot showing detection of various higher molecular weight (HMW) species of TRIM37 upon treatment with increasing concentrations of DSS crosslinker. Vinculin is used as a loading and oligomerization control. Dotted lines indicate separate cropped sections of the same immunoblot. Representative data; *n* = 3 biological replicates. Bottom, Densitometric analysis of immunoblot with a graph depicting normalized HMW TRIM37 intensity upon increasing DSS concentrations relative to DMSO control (−DSS). Mean ± s.e.m. (D) Representative Sanger sequencing traces of the *TRIM37* locus in parental and CRISPR–Cas9 edited RPE-1 *TRIM37^C18R^* cells, highlighting the mutation (TGT>CGT) responsible for the biallelic C18R residue substitution, denoted by an asterisk. (E) Left, immunoblot showing endogenous TRIM37 protein levels across the indicated cellular fractions from RPE-1 *TRIM37^C18R^*cells. Validation markers include CEP192, Centrobin, and SAS6 for centrosomal proteins, and Lamin A/C for the nuclear fraction. Ponceau-stained blot indicates loading. Representative data; *n* = 3 biological replicates. WCE, whole-cell extract; exp, exposure. Right, Densitometric analysis of immunoblot in with a graph depicting endogenous TRIM37 enrichment in indicated fractions relative to WCE. *P* values, one-way ANOVA with post hoc Dunnett’s multiple comparisons test to evaluate enrichment of TRIM37 in each cellular fraction relative to WCE. Mean ± s.e.m. (F) Left, immunoblot showing detection of various higher molecular weight (HMW) species of endogenous TRIM37 upon treatment of RPE-1 *TRIM37^C18R^* cells with increasing concentrations of DSG crosslinker. Vinculin is used as a loading and oligomerization control. Representative data; *n* = 3 biological replicates. Right, Densitometric analysis of immunoblot with a graph depicting normalized HMW TRIM37 intensity upon increasing DSG concentrations relative to DMSO control (−DSG). Mean ± s.e.m.

**Extended Data Figure 3.**
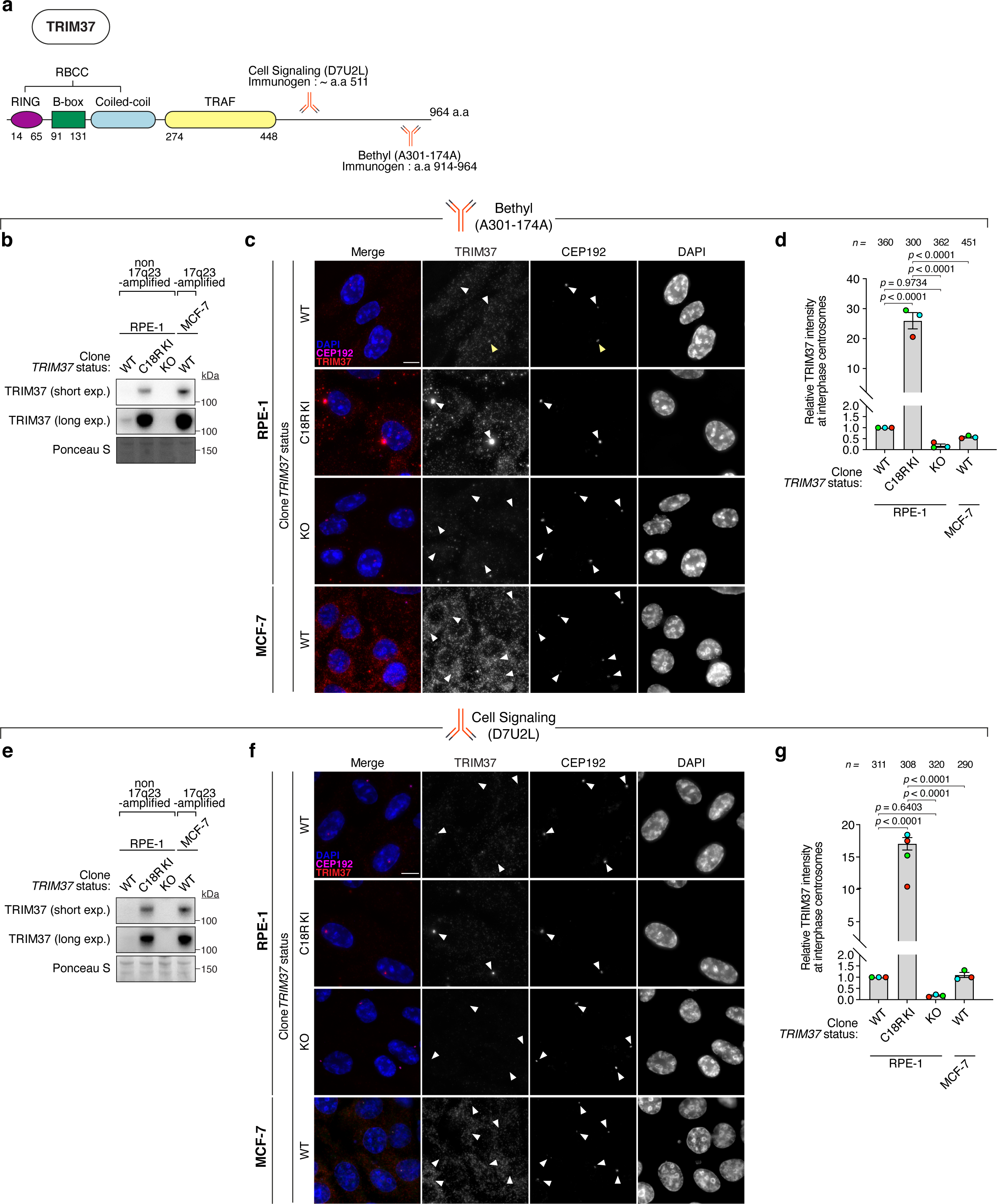
Endogenous TRIM37 localization at the centrosome is revealed by E3 ligase inactivation. (A) Schematic of the TRIM37 protein, highlighting epitopes recognized by two distinct commercial antibodies. (B-D) The commercial TRIM37 antibody (Bethyl, A301-173A) was utilized for the following experiments. (B) Immunoblot showing endogenous TRIM37 protein levels across a panel of cell lines with the indicated *TRIM37* status. Ponceau-stained blot indicates loading. Representative data; *n* = 3 biological replicates. KI, knock-in; KO, knock-out; exp, exposure. (C) Representative images showing the immunostaining pattern of endogenous TRIM37 in the cell line panel. Arrows indicate the location of centrosomes. Representative data; *n* = 3 biological replicates. Scale bars, 5 μm. (D) Quantification of endogenous TRIM37 signal at the centrosomes of the cell line panel. *n* = 3 biological replicates, each with >100 cells. *P* values, one-way ANOVA with post hoc Tukey’s multiple comparisons test. Mean ± s.e.m. (E-G) Same as in (B-D), but with a second commercial antibody (Cell Signaling Technology, D7U2L).

**Extended Data Figure 4.**
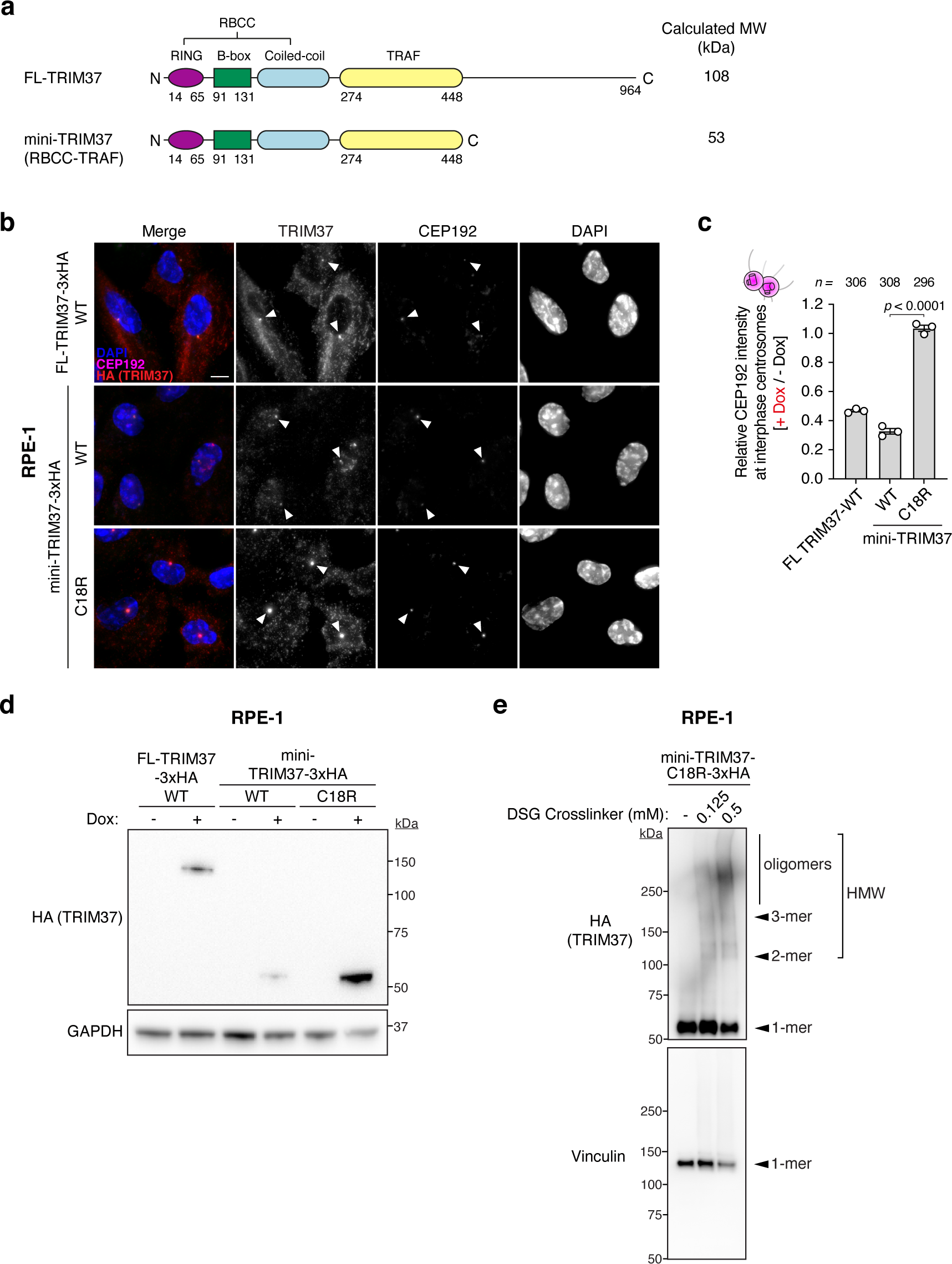
Defining the minimal TRIM37 domain architecture required for centrosome regulation. (A) Schematic of the miniTRIM37 (RBCC-TRAF) construct compared to full-length TRIM37. (B) Representative images of the localization and effect of indicated HA-tagged TRIM37 constructs on centrosomal CEP192 levels in RPE-1 tet-on TRIM37 cells. Arrows indicate the location of centrosomes. Representative data; *n* = 3 biological replicates. Scale bars, 5 μm. (C) Quantification of centrosomal CEP192 signal upon doxycycline-induced expression of indicated HA-tagged TRIM37 constructs in RPE-1 tet-on TRIM37 cells from (B). *n* = 3 biological replicates, each with >100 cells. *P* values, one-way ANOVA with post hoc Tukey’s multiple comparisons test. Mean ± s.e.m. (D) Immunoblot showing total protein levels of indicated HA-tagged TRIM37 constructs in RPE-1 tet-on TRIM37 cells from (B-C). GAPDH, loading control. Representative data; *n* = 3 biological replicates. (E) Immunoblot showing detection of various higher molecular weight (HMW) species of miniTRIM37 upon treatment with increasing concentrations of DSG crosslinker. Vinculin is used as a loading and oligomerization control. Representative data; *n* = 3 biological replicates.

**Extended Data Figure 5.**
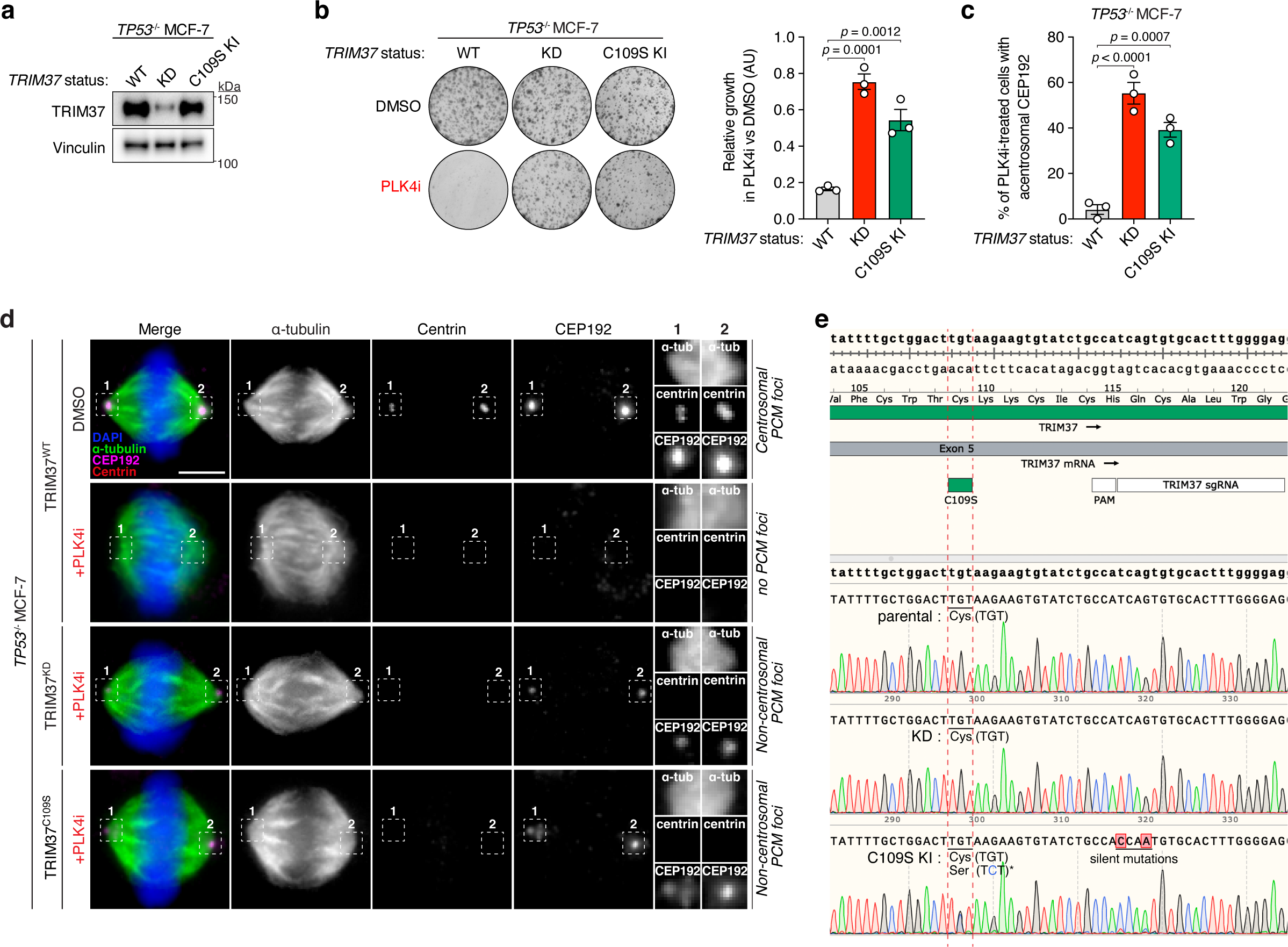
Impairment of TRIM37 oligomerization attenuates synthetic lethality in 17q23-amplified cells with PLK4 inhibition. (A) Immunoblot showing TRIM37 protein levels in *TP53*^−/−^ MCF-7 cells. TRIM37 wild-type (WT), *TRIM37* knockdown (KD) via shRNA, and cells harboring the C109S mutation in approximately half of the TRIM37 alleles present (*TRIM37*^C109S^) were used. Vinculin, loading control. Representative data; *n* = 3 biological replicates. (B) Left, Representative data of a 10-d clonogenic survival of indicated MCF-7 cell lines from (A) treated with DMSO (control) or PLK4 inhibitor (PLK4i) (125 nM). Right, Quantification of relative growth in the presence PLK4i relative to DMSO. *P* values, one-way ANOVA with post hoc Dunnett’s multiple comparisons test to evaluate differences between each experimental condition (KD and C109S) and WT. Mean ± s.e.m (C) Quantification of mitotic CEP192 foci in PLK4i-treated *TP53*^−/−^ MCF-7 cells that lack centrosomes. *n* = 3, biological replicates, each comprising >30 cells. *P* values, one-way ANOVA with post hoc Dunnett’s multiple comparisons test to evaluate differences between each experimental condition (KD and C109S) and WT. Mean ± s.e.m (D) Representative images for (C). Scale bars, 5 μm. (E) Representative Sanger sequencing traces for the *TRIM37* locus in parental *TP53*^−/−^ MCF-7 cells subjected to *TRIM37* knockdown (KD) via shRNA, and CRISPR–Cas9 edited *TRIM37^C109S^* KI cells. The mutation (TGT>TCT) leading to the C109S residue substitution is denoted by an asterisk. Silent mutations introduced to prevent re-editing are highlighted.

**Extended Data Figure 6.**
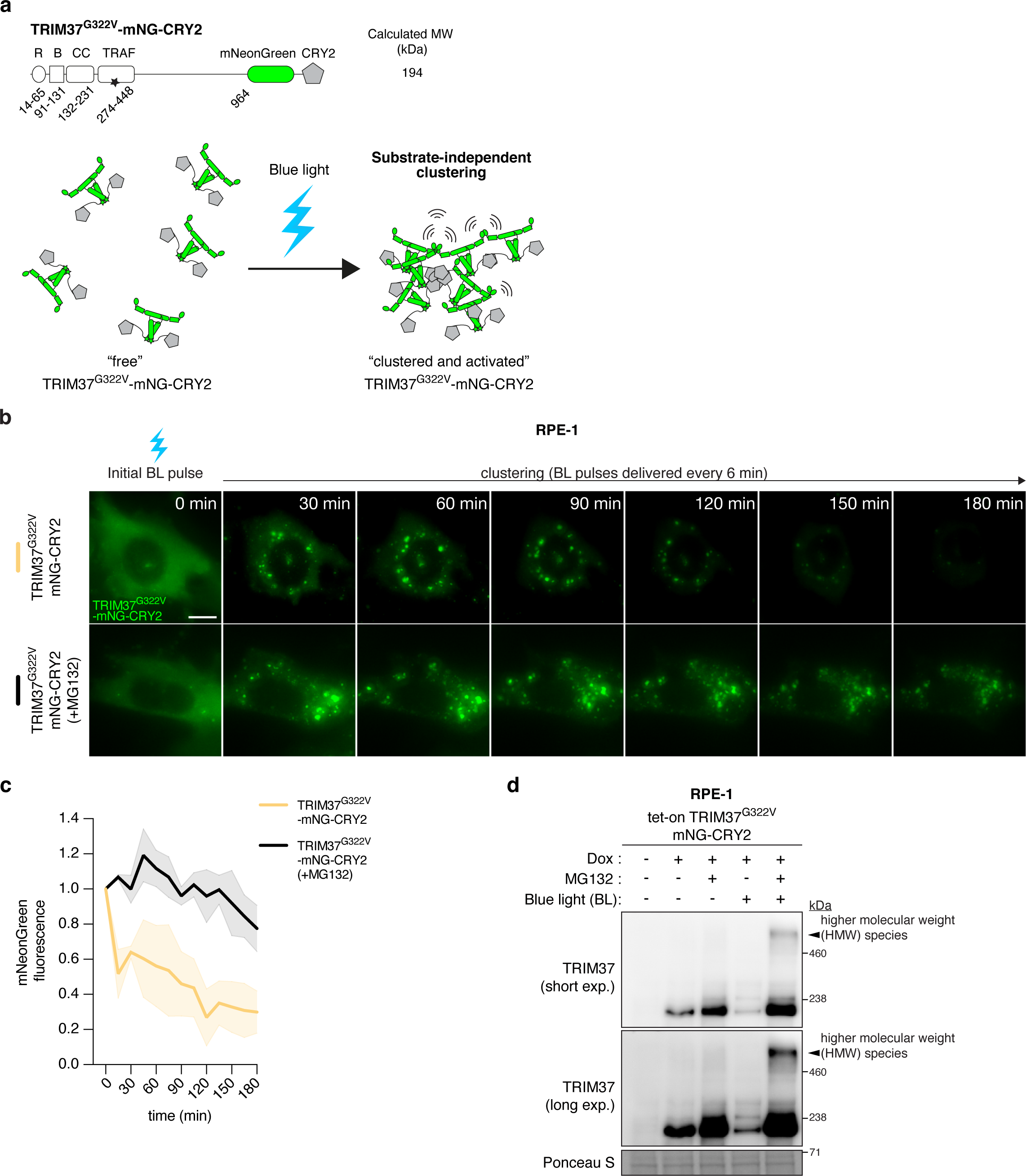
Substrate-independent clustering is sufficient to activate TRIM37. (related to Figure 6). (A) Top, schematic of the TRIM37^G322V^-mNeonGreen-CRY2 optogenetic fusion construct. The star denotes the TRAF domain mutation (G322V). Bottom, illustration of the blue light (BL)-activated optogenetic system enabling TRIM37 clustering independent of binding to a centrosome substrate. (B) Representative time-lapse images of RPE-1 cells expressing the optogenetic construct detailed in (A) incubated in the presence or absence of MG132. Timestamps indicate minutes post blue light exposure. Scale bar = 10 µm. (C) Quantification of mNeonGreen fluorescence intensity from (B), with each condition comprising >30 cells. Mean ± s.d. (D) RPE-1 cells expressing optogenetic constructs detailed in (A) were incubated with or without doxycycline (Dox) and MG132 (10 μM) in the absence or presence blue light for 3 h before immunoblotting for the indicated proteins. Higher molecular weight (HMW) TRIM37 species were prominently formed only in MG132 and BL-stimulated conditions and are indicated with an arrow. Ponceau-stained blot indicates loading. Representative data; *n* = 3 biological replicates. exp, exposure.

